# A plant receptor-like kinase promotes cell-to-cell spread of RNAi and is targeted by a virus

**DOI:** 10.1101/180380

**Authors:** Tabata Rosas-Diaz, Dan Zhang, Pengfei Fan, Liping Wang, Xue Ding, Yuli Jiang, Tamara Jimenez-Gongora, Laura Medina-Puche, Xinyan Zhao, Zhengyan Feng, Guiping Zhang, Xiaokun Liu, Eduardo R Bejarano, Li Tan, Jian-Kang Zhu, Weiman Xing, Christine Faulkner, Shingo Nagawa, Rosa Lozano-Duran

**Affiliations:** Shanghai Center for Plant Stress Biology, Chinese Academy of Sciences, Shanghai 201602, China; University of the Chinese Academy of Sciences, Beijing 100049, China; John Innes Centre, Norwich, United Kingdom; University of Malaga, Malaga, Spain; Department of Horticulture and Landscape Architecture, Purdue University, West Lafayette, 47907, IN, USA; CAS-JIC Centre of Excellence for Plant and Microbial Science (CEPAMS), Shanghai Institutes for Biological Sciences, Chinese Academy of Sciences (CAS), Shanghai 200032, China

## Abstract

RNA interference (RNAi) in plants can move from cell to cell, allowing for systemic spread of an anti-viral immune response. How this cell-to-cell spread of silencing is regulated is currently unknown. Here, we describe that the C4 protein from *Tomato yellow leaf curl virus* has the ability to inhibit the intercellular spread of RNAi. Using this viral protein as a probe, we have identified the receptor-like kinase (RLK) BARELY ANY MERISTEM 1 (BAM1) as a positive regulator of the cell-to-cell movement of RNAi, and determined that BAM1 and its closest homologue, BAM2, play a redundant role in this process. C4 interacts with the intracellular domain of BAM1 and BAM2 at the plasma membrane and plasmodesmata, the cytoplasmic connections between plant cells, interfering with the function of these RLKs in the cell-to-cell spread of RNAi. Our results identify BAM1 as an element required for the cell-to-cell spread of RNAi and highlight that signalling components have been co-opted to play multiple functions in plants.

## MAIN TEXT

RNA interference (RNAi) mediated by small interfering RNA (siRNA) is considered the main anti-viral defence mechanism in plants. RNAi relies on the production of virus-derived siRNA (vsiRNA) by RNaseIII Dicer-like proteins, mainly DCL2 and its surrogate DCL4; vsiRNAs are then loaded into argonaute (AGO) proteins AGO1- and AGO2-containing complexes to target viral RNA for cleavage (*1*). siRNAs can move from cell to cell (*2*), presumably symplastically through plasmodesmata (PD), and it has been proposed that vsiRNAs can move ahead of the front of the infection, leading to spread of silencing and immunizing plant tissues prior to the arrival of the virus. However, how the cell-to-cell movement of siRNAs/vsiRNAs occurs remains elusive, and efforts to identify plant proteins specifically involved in this process through forward genetics have not been fruitful (*3*, 4).

In order to counter anti-viral RNAi, viruses have evolved viral suppressors of RNA silencing (VSR). All plant viruses described to date encode at least one VSR, supporting the notion that active suppression of RNAi is a *conditio sine qua non* for infectivity. Although independently evolved VSRs have been shown to target different steps of the RNAi pathway (*5*), direct interference of a VSR with cell-to-cell movement of the silencing signal has not yet been characterized.

The C4 protein from *Tomato yellow leaf curl virus* (TYLCV; family *Geminiviridae*) can delay the systemic spread of silencing in transgenic *Nicotiana benthamiana* plants, but is not a local suppressor of RNAi (*6*). We therefore set out to assess whether C4 could interfere with cell-to-cell movement of silencing. Interestingly, we found that C4 localizes to different compartments in the plant cell, mainly plasma membrane (PM), plasmodesmata (PD), and chloroplasts; PM/PD localization depends on the presence of a myristoylation motif at the N-terminus of the protein (Figure 1; Figure S1). A mutant virus expressing a non-myristoylable version of C4 (C4_G2A_), which localizes to chloroplasts exclusively, displays severely compromised infectivity in tomato (Figure S2), suggesting that PM/PD localization is required for full infectivity. When expressed constitutively in Arabidopsis, C4 causes strong developmental alterations, which also requires myristoylation (Figure 1, Figure S3).

**Figure 1.**
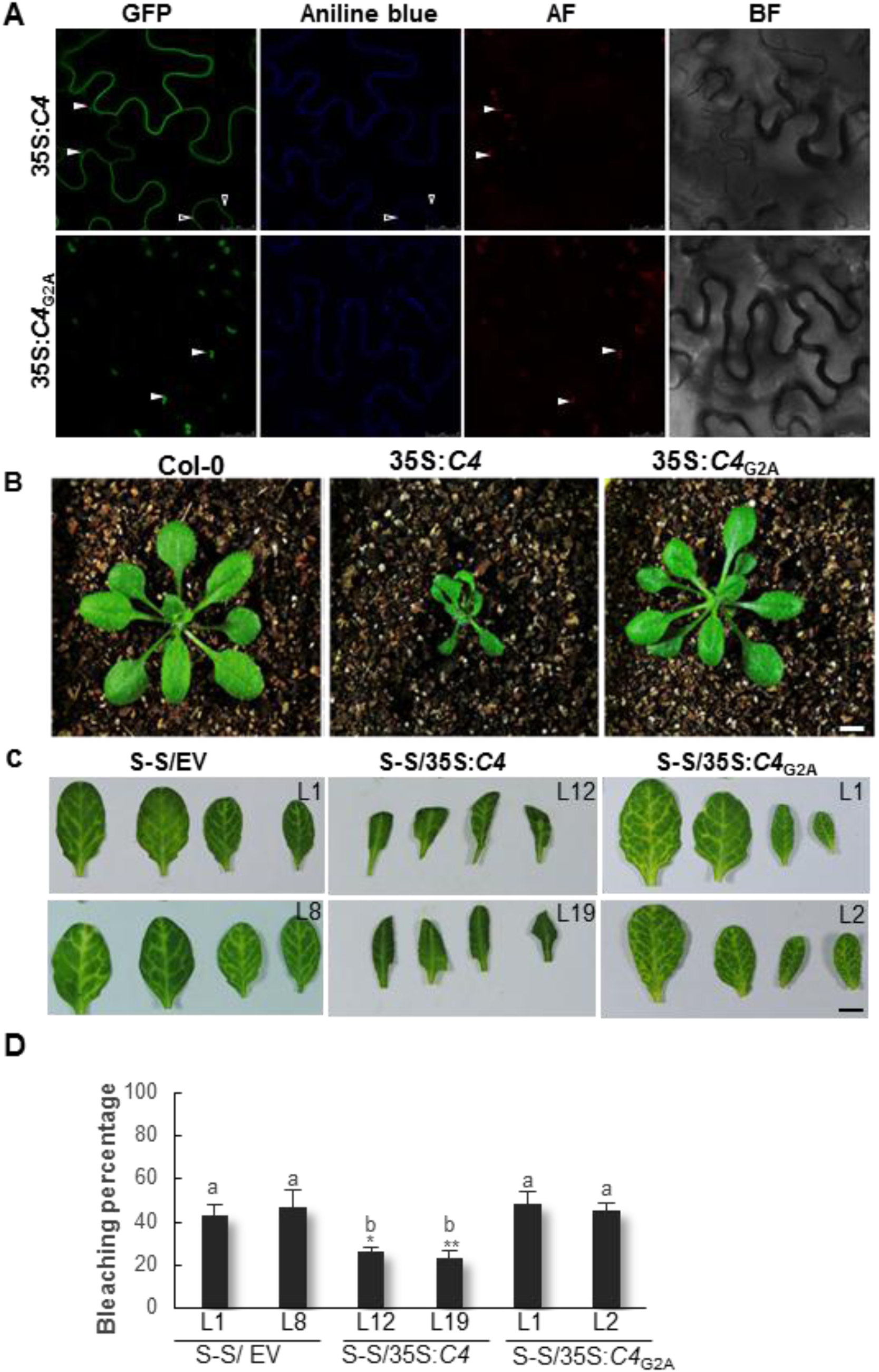
Plasma membrane/plasmodesmal C4 suppresses cell-to-cell spread of RNAi. A. Subcellular localization of C4-GFP or the non-myristoylable mutant C4_G2A_-GFP upon transient expression in *Nicotiana benthamiana* leaves. AF: Autofluorescence. BF: Bright field. Empty arrowheads indicate plasmodesmata; filled arrowheads indicate chloroplasts. B. Transgenic plants expressing C4 or the non-myristoylable mutant C4_G2A_ at five weeks post-germination. C. Leaves of four-week-old transgenic SUC:*SUL* plants expressing C4 (S-S/35S:*C4*), the non-myristoylable mutant C4_G2A_ (S-S/35S:*C4*_*G2A*_), or transformed with the empty vector (S-S/EV). Each set of four leaves comes from one T2 plant from an independent line. D. Quantification of the bleaching percentage of the leaves in (B). Bars represent SE; bars with the same letter are not significantly different (P=0.05) according to Dunnet’s multiple comparison test. Scale bar: 0.5 cm.

To investigate whether C4 affects the cell-to-cell spread of RNAi, we expressed this protein in the SUC:*SUL* silencing reporter system (*7*). SUC:*SUL* transgenic Arabidopsis plants express an inverted repeat of the endogenous *SULPHUR* (*SUL*) gene (At4g18480) in phloem companion cells, which leads to the production of 21 and 24 nt siRNAs against *SUL* and the ensuing RNAi, resulting in a chlorotic phenotype of the silenced cells. Cell-to-cell movement of 21 nt siRNAs causes the spread of silencing to 10-15 cells beyond the vasculature ((*3*); Figure 1). Expression of wild type C4, but not of the non-myristoylable mutant C4_G2A_, greatly diminishes the spread of the silencing phenotype, suggesting that PM/PD-localized C4 may interfere with the cell-to-cell movement of silencing from the vasculature (Figure 1, Figure S4). Of note, expression of C4 triggered a detectable decrease in the accumulation of siRNAs against *SUL* (Figure S10), indicating that this viral protein might act as a VSR at several different levels.

With the aim of uncovering the molecular mechanism of the C4-mediated suppression of cell-to-cell spread of silencing, we searched for interacting partners of C4 through yeast two-hybrid screening of a TYLCV-infected tomato cDNA library, and affinity purification followed by mass spectrometry analyses (AP-MS) upon transient expression of C4-GFP in *N. benthamiana*. Interestingly, these independent approaches both identified the receptor-like kinase (RLK) BARELY ANY MERISTEM 1 (BAM1) as an interactor of C4; the interaction with C4 occurs through the kinase domain of BAM1, as shown by mapping of this interaction in yeast (Figure 2). The interaction between C4 and BAM1 from Arabidopsis was confirmed using co-immunoprecipitation (Co-IP), Förster resonance energy transfer – fluorescence lifetime imaging (FRET-FLIM), bimolecular fluorescence complementation (BiFC), and gel filtration chromatography (Figure 2), and it requires PM/PD localization of C4 (Figure S5). The interaction of C4 with BAM1 from tomato, natural host of TYLCV, was confirmed by BiFC and FRET-FLIM (Figure S6). Interestingly, *BAM1* is strongly expressed in the vasculature, to which many viruses, including TYLCV, are restricted ((*8*); Figure S7). BiFC assays revealed that C4 interacts with BAM1 at PM and PD (Figure 2), where both proteins co-localize (Figure S8), thus it is conceivable that BAM1 participates in the cell-to-cell movement of silencing and is targeted by C4. In order to test if BAM1 is indeed involved in this process, we generated transgenic SUC:*SUL* plants overexpressing tagged and untagged versions of the endogenous protein (SUC:*SUL*/35S:*BAM1*, SUC:*SUL*/35S:*BAM1-GFP*, SUC:*SUL*/35S:*BAM1-FLAG*) as well as tomato BAM1 (SUC-SUL/35S:*SlBAM1-GFP)*. In all cases, overexpression of *BAM1* resulted in an extended spread of silencing from the vasculature (Figure 3; Figures S6, S9, S10) while accumulation of siRNA against *SUL* remained unaltered (Figure S10), indicating that BAM1 promotes cell-to-cell movement of the silencing signal. Kinase activity does not seem to be required for this function, since overexpression of a kinase-dead BAM1_D820N_ mutant, mutated in the catalytic aspartic acid, produces a promotion of the spread of silencing indistinguishable to that of the wild type protein (Figure S11). A mutation in the *BAM1* gene alone did not affect the silencing phenotype of the SUC:*SUL* plants (Figure 3), which suggests functional redundancy. In Arabidopsis, *BAM1* has two close homologues, named *BAM2* and *BAM3*; the kinase domains of BAM1 and BAM2 are 94% identical at the protein level (Figure S13), whereas those of BAM1 and BAM3 are 73% identical. Consistent with this, C4 can also interact with BAM2 and, more weakly, with BAM3, as observed in co-IP, BiFC, and FRET-FLIM assays (Figure 4). Strikingly, simultaneous mutation of *BAM1* and *BAM2* in SUC:*SUL* plants by CRISPR-Cas9-mediated genome editing (*9*) (Figure S14; Table S1) markedly diminished the spread of silencing from the vasculature, strongly resembling the effect of C4 expression (Figure 4). Similar to C4 transgenic plants, silencing of *SUL* can still be observed around the mid vein in *bam1 bam2*/SUC:*SUL* plants, but spread of silencing from secondary veins is basically abolished. Therefore, BAM1 and BAM2 redundantly promote cell-to-cell movement of silencing from the vasculature, and C4 targets both homologues to inhibit their activity in this process. Although other functions of BAM1/BAM2 may also be inhibited by C4, as suggested by similarities in developmental defects of *bam1 bam2* mutants and C4 transgenic plants ((*10*); Jian et al., in preparation; Figure S15), C4 does not impair all activities of BAM1, since C4 transgenic plants display a wild type-like response to CLV3p in root elongation assays, which is BAM1-dependent (*11*) (Figure S16). Taken together, our results show that C4 targets BAM1/BAM2 and suppresses a subset of the functions of these RLKs, including the promotion of the cell-to-cell spread of silencing from the vasculature. This effect of BAM1/BAM2 on intercellular spread of RNAi is specific, since general plasmodesmal conductance is not affected by expression of *C4* or overexpression of *BAM1* (Figure S17).

**Figure 3.**
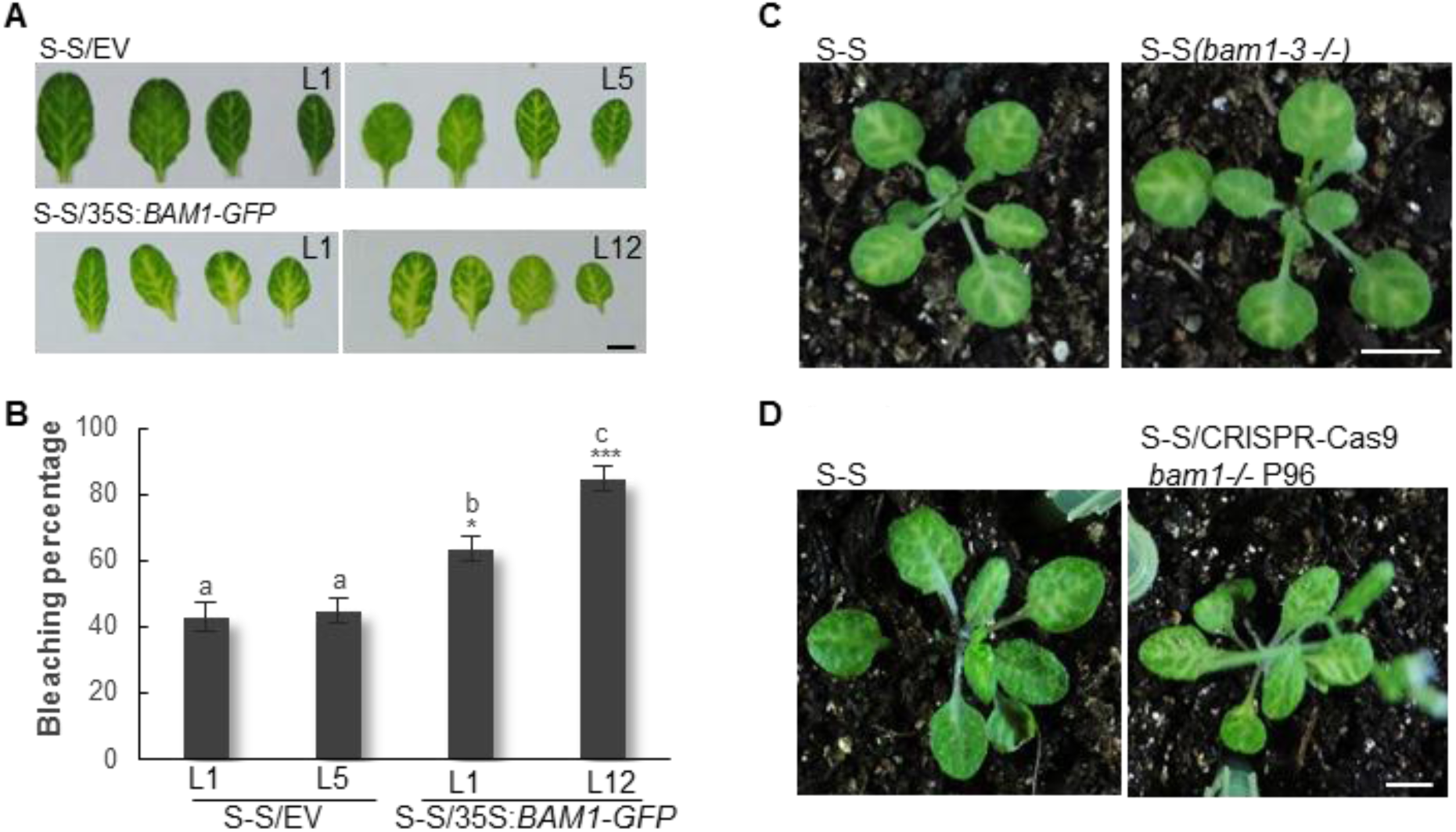
BAM1 promotes the cell-to-cell spread of RNAi. A. Leaves of four-week-old transgenic SUC:*SUL* plants overexpressing BAM1-GFP (S-S/35S:*BAM1-GFP*) or transformed with the empty vector (S-S/EV). Each set of leaves comes from one T2 plant from an independent line. B. Quantification of the bleaching percentage of the leaves in (A). Bars represent SE; bars with the same letters are not significantly different (P=0.05) according to Dunnet’s multiple comparison test. C and D. Phenotype of SUC:*SUL* plants mutated in *bam1*. (C) depicts a *bam1-3* mutant homozygous for the SUC:*SUL* transgene (F3); (D) depicts a *bam1* mutant obtained by genome editing with CRISPR-Cas9 (for details, see Figure S12 and Table S1). Scale bar: 0.5 cm.

**Figure 4.**
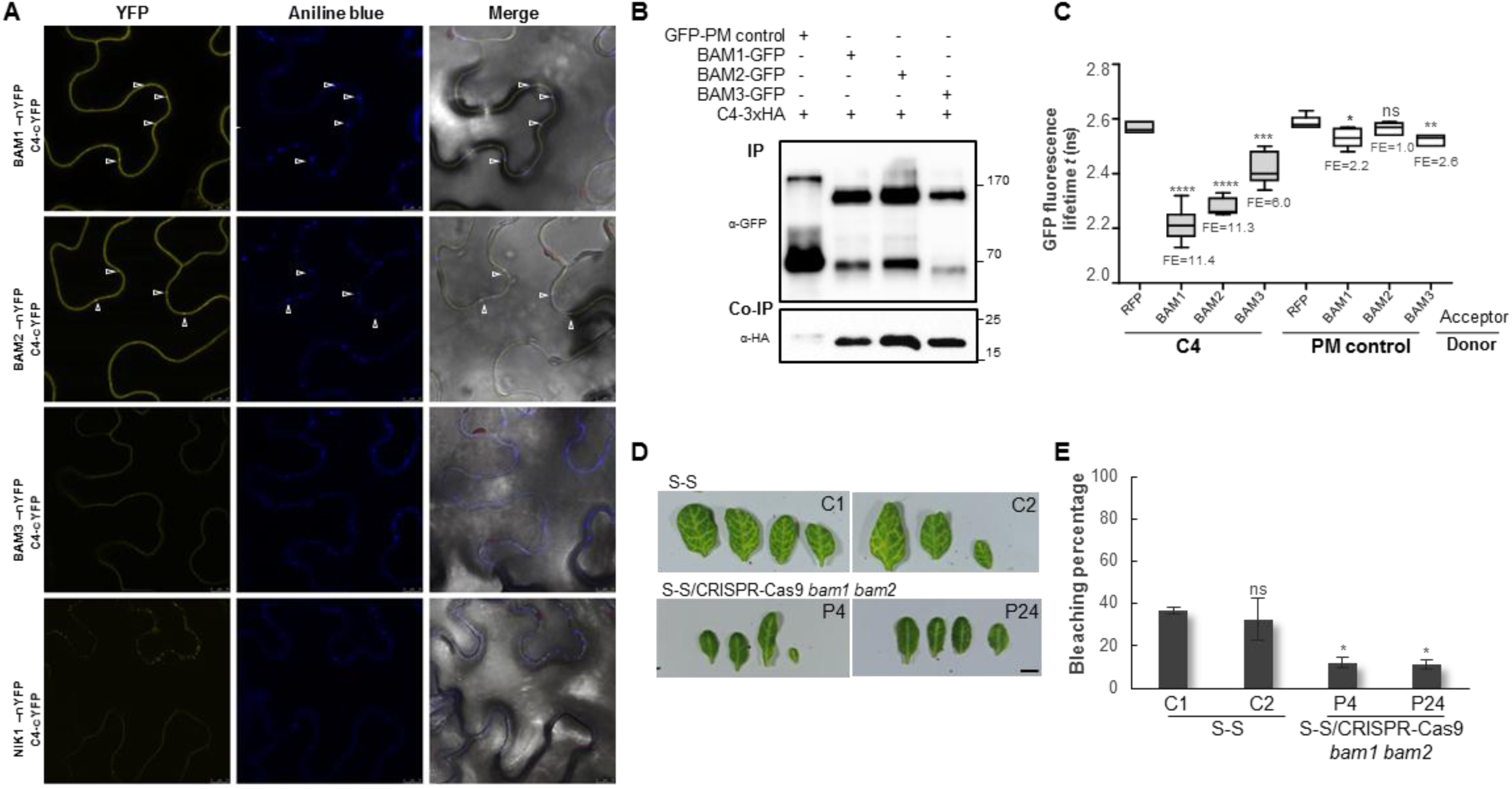
BAM1 and its homologue BAM2 are required for the cell-to-cell spread of RNAi. A. Interaction between C4 and BAM1 and C4 and BAM2 by BiFC upon transient co-expression in *N. benthamiana* leaves. The receptor-like kinase NIK1 is used as negative control. Arrowheads indicate plasmodesmata. B. Co-immunoprecipitation of BAM1-GFP, BAM2-GFP, and BAM3-GFP with C4-3xHA upon transient co-expression in *N. benthamiana* leaves. Numbers on the right indicate molecular weight. C. Interaction between C4 and BAM1, BAM2, and BAM3 by FRET-FLIM upon transient co-expression in *N. benthamiana* leaves. The membrane protein NP_564431 (NCBI) is used as a negative control (PM control). FE: FRET efficiency. Asterisks indicate a statistically significant difference (****,p-value < 0.0001; ***,p-value < 0.005; **, p-value < 0.01; *, p-value < 0.05), according to a Student’s t-test; ns: not significant. D. Leaves of four-week-old transgenic SUC:*SUL* plants mutated in *bam1* and *bam2* obtained by genome editing with CRISPR-Cas9 (for details, see Figure S14 and Table S1). Each set of leaves comes from one T3 plant from an independent line. Scale bar: 0.5 cm. E. Quantification of the bleaching percentage of the leaves in (D). Bars represent SE. Asterisks indicate a statistically significant difference (*, p-value < 0.05), according to a Student’s t-test; ns: not significant.

**Figure 2.**
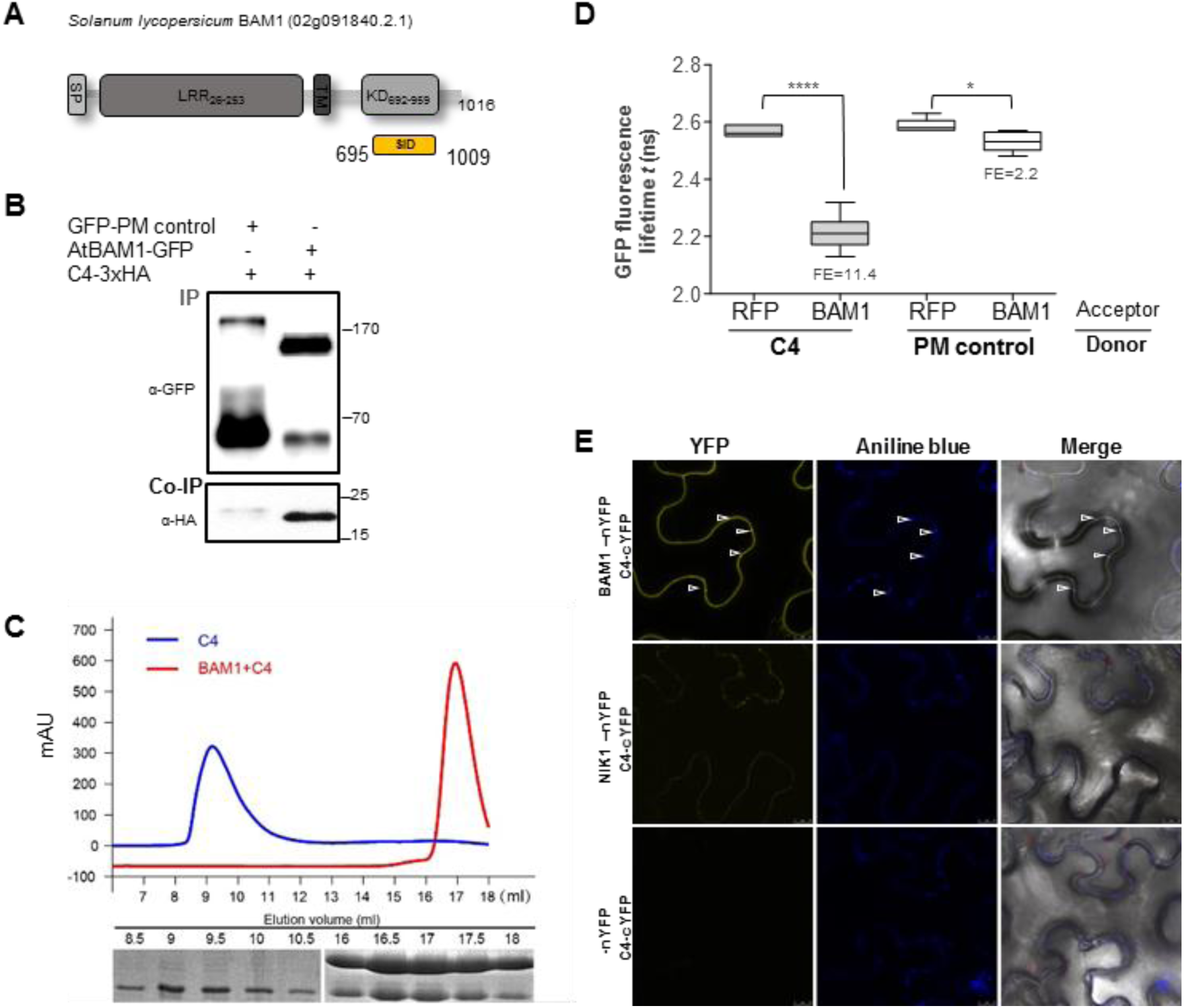
C4 interacts with the receptor-like kinase BAM1. A. Schematic representation of the receptor-like kinase BAM1 from tomato (*Solanum lycopersicum*) isolated in the yeast two-hybrid screen as an interactor of C4. The signal peptide (SP), leucine-rich repeats (LRR), transmembrane domain (TM), and kinase domain (KD) are shown; numbers indicate the beginning and end of each domain, in amino acids. The selected interaction domain (SID) is the minimal fragment found to interact with C4 in this screen. B. Co-immunoprecipitation of BAM1-GFP from Arabidopsis with C4-3xHA upon transient co-expression in *N. benthamiana* leaves. Numbers on the right indicate molecular weight. C. Gel filtration chromatography of C4 and the C4/BAM1 complex expressed in *Escherichia coli*. Top panel: gel filtration chromatograms of C4 and the C4/BAM1 complex. Bottom panel: Coomassie blue staining of the peak fractions shown above following SDS-PAGE. The elution volume of C4 shifted from 9mL to 17mL after co-expression with BAM1, indicating an interaction between these two proteins. D. Interaction between C4 and BAM1 by FRET-FLIM upon transient co-expression in *N. benthamiana* leaves. The membrane protein NP_564431 (NCBI) is used as a negative control (PM control). FE: FRET efficiency. Asterisks indicate a statistically significant difference (****,p-value < 0.0001; *, p-value < 0.05), according to a Student’s t-test. E. Interaction between C4 and BAM1 by BiFC upon transient co-expression in *N. benthamiana* leaves. The receptor-like kinase NIK1 is used as a negative control. Arrowheads indicate plasmodesmata.

Using a viral protein as a probe, we have identified the PM/PD-localized RLK BAM1 as a positive regulator of the cell-to-cell movement of RNAi from the vasculature, and determined that BAM1 and BAM2 are functionally redundant in this process. Since pathogen effectors have evolved to overcome genetic redundancy, their use to investigate novel cellular processes is a valuable complement to traditional forward genetic screens.

BAM1/BAM2 had been previously characterized with respect to their role in development (*11-13*). Our work identifies a novel function of these RLKs in the cell-to-cell spread of RNAi; however, the underlying molecular mechanism is still elusive. The finding that kinase activity seems to be expendable for the BAM1-mediated promotion of cell-to-cell spread of silencing from the vasculature suggests that these RLKs may have a scaffolding rather than an enzymatic function in this process. Whether BAM1/BAM2 can bind siRNA-binding proteins, such as AGO proteins, or siRNA molecules directly remains to be determined. Strikingly, C4 seems to inhibit certain functions of BAM1/BAM2, including the promotion of intercellular RNAi spread, but not all: this differential targeting may have evolved to avoid detrimental pleiotropic effects of unspecifically suppressing every role of a multifunctional target. The specificity of the C4-mediated inhibition of BAM1 functions may be determined by subcellular localization and composition of the potential BAM1-containing protein complexes involved in the different signalling pathways.

Another open question is whether BAM1/BAM2 also participate in the cell-to-cell movement of other sRNAs, such as miRNAs or tasiRNAs, and if C4 targets these non-cell-autonomous processes as well, dependent or independently of BAM1/BAM2. Mobile sRNAs determine the proper development of xylem and establishment of leaf polarity (*14-17*); the potential role of BAM1/BAM2 and the effect of C4 in these processes is currently under investigation.

## ACKNOWLEDGEMENTS

The authors thank Tsuyoshi Nakagawa, Zachary Nimchuk, Yuanzheng Wang, and Alberto Macho for kindly sharing materials; the Proteomics facility at the Shanghai Center for Plant Stress Biology; Wenjie Zeng, Xinyu Jian, Aurora Luque, and Yujing (Ada) Liu for technical assistance; Sebastian Wolf for critical reading of the manuscript; and all members in Rosa Lozano-Duran’s and Alberto Macho’s groups for stimulating discussions and helpful suggestions. This research was supported by the Centre of Excellence for Plant and Microbial Sciences (CEPAMS), established between the John Innes Centre and the Chinese Academy of Sciences and funded by the UK Biotechnology and Biological Sciences Research Council and the Chinese Academy of Sciences, and a Mianshang grant from the National Natural Science Foundation of China (NSFC) to RL-D (grant number 31671994). Research in RL-D’s lab is funded by the Shanghai Center for Plant Stress Biology of the Chinese Academy of Sciences and the 100 Talent program of the Chinese Academy of Sciences. TR-D is the recipient of a President’s International Fellowship Initiative (PIFI) postdoctoral fellowship (No. 2016PB042) from the Chinese Academy of Sciences. TJ-G is sponsored by a CAS-TWAS President’s Fellowship for International PhD students.

## AUTHOR CONTRIBUTIONS

TR-D, DZ, PF, LW, XD, YJ, TJ-G, LM-P, XZ, ZF, GZ, and XL performed experiments and analysed data. TR-D, ERB, LT, J-KZ, WX, CF, SN, and RL-D designed, supervised, and interpreted experiments. CF and RL-D conceived the original project. RL-D wrote the manuscript, with input from all authors.

## COMPETING FINANCIAL INTERESTS

The authors declare no competing financial interests.

## MATERIALS AND CORRESPONDENCE

Correspondence and material requests should be addressed to Rosa Lozano-Duran.

## SUPPLEMENTARY MATERIAL

## MATERIALS AND METHODS

### Plant material

All Arabidopsis mutants and transgenic plants used in this work are in the Col-0 background. The *bam1-3* mutant is described in (*18*). The SUC:*SUL* transgenic line is described in (*19*). To generate the *C4*, *C4*_*G2A*_, pBAM1:*YFP-NLS*, and 35S:BAM1 lines, wild type Arabidopsis plants were transformed with pGWB2-*C4*, pGWB2-*C4*_*G2A*_,, pBAM1:*YFP-NLS*, and pGWB2-*BAM1*, respectively (see Plasmids and cloning). To generate the SUC:*SUL*/*C4*, SUC:*SUL*/*C4*_*G2A*_, SUC:*SUL*/35S:*BAM1*, SUC:*SUL*/35S:*BAM1-GFP*, SUC:*SUL*/35S:*BAM1-FLAG*, and SUC:*SUL*/35S:*SlBAM1-GFP* lines, SUC:*SUL* plants were transformed with the corresponding plasmids (pGWB2-*C4*, pGWB2-*C4*_*G2A*_, pGWB2-*BAM1*, pGWB505-*BAM1*, pGWB511-*BAM1*, and pGWB505-*SlBAM1*, respectively) (see Plasmids and cloning) using the floral dipping method. To generate the *bam1-3*/SUC:*SUL* plants, SUC:*SUL* plants were crossed to *bam1-3* plants, and homozygous individuals for both mutation and transgene were identified and phenotyped in F3. To generate the *bam1 bam2*/SUC:*SUL* lines, SUC:*SUL* plants were transformed with CRISPR-Cas9 constructs to target *BAM1* and *BAM2* (see Plasmids and cloning). Mutation of *BAM1* and *BAM2* was confirmed in T1 by sequencing, and those plants carrying mutations were selected for further characterization in subsequent generations. Homozygous or biallelic mutant plants were identified by sequencing and phenotyped in T2 and T3.

The tomato cultivar used in this work for viral infection assays and cloning of *SlBAM1* is Money maker.

### Co-immunoprecipitation and protein analysis

Protein extraction, co-immunoprecipitation, and proteins analysis were performed as described in (*20*), with minor modifications. In order to efficiently extract membrane proteins, 2% of NP-40 was used in the protein extraction buffer. The antibodies used are as follows: anti-GFP (Abiocode M0802-3a), anti-HA (Santa-Cruz sc-7392), anti-Rabbit IgG (Sigma A0545), and anti-Mouse IgG (Sigma A2554).

### Confocal imaging

*N. benthamiana* plants were agroinfiltrated with clones to express C4-GFP, C4-RFP, C4_G2A_-GFP, and BAM1-GFP, and samples were imaged two days later on a Leica TCS SP8 point scanning confocal microscope using the pre-set settings for GFP with Ex:489nm, Em:500-500nm, and for RFP with Ex:554, Em:580-630. For callose deposition at plasmodesmata, samples were stained with a solution of 0.05% (w/v) aniline blue in water by infiltration into leaves 30 minutes before imaging with Ex:405 nm, Em: 448-525 nm.

### Bimolecular fluorescent complementation

The plasmids used for BiFC are described in (*21*) (see Plasmids and cloning). *N. benthamiana* plants were agroinfiltrated with clones to express the corresponding proteins, and samples were imaged two days later on a Leica TCS SP8 confocal microscope, using the pre-set settings for YFP with Ex:514 nm, Em: 525-575 nm.

### FRET-FLIM imaging

For FRET-FLIM experiments, donor proteins (fused to GFP) were expressed from vectors pGWB5 or pGWB606, and acceptor proteins (fused to RFP) were expressed from vector pB7WRG2.0. FRET-FLIM experiments were performed on a Leica TCS SMD FLCS confocal microscope excitation with WLL (white light laser) and emission collected by a SMD SPAD (single photon sensitive avalanche photodiodes) detector. Leaf discs of *N. benthamiana* plants transiently co-expressing donor and acceptor, as indicated in the figures, were visualized two days after agroinfiltration. Accumulation of the GFP- and RFP-tagged proteins was estimated prior to measuring lifetime. The tunable WLL set at 489 nm with a pulsed frequency of 40 MHz was used for excitation, and emission was detected using SMD GFP/RFP Filter Cube (with GFP:500–550 nm). The fluorescence lifetime shown in the figures corresponding to the average fluorescence lifetime of the donor (τ) was collected and analysed by QicoQuant SymphoTime software. Lifetime is normally amplitude weighted mean value using the data from the single (GFP fused donor protein only or GFP fused donor protein with free RFP acceptor or with non-interacting RFP fused acceptor protein) or bi-exponential fit (GFP fused donor protein interacting with RFP fused acceptor protein). Mean lifetimes are presented as means ± SD based on more than 10 cells from at least three independent experiments. FRET efficiency was calculated according to the formula E = 1-τDA/τD, where τDA is the average lifetime of the donor in the presence of the acceptor and τD is the average lifetime of the donor in the absence of the acceptor.

### Plasmids and cloning

Plasmids and primers used for cloning are summarized in Table S2. The TYLCV clone used as template is AJ489258 (GenBank).

Vectors from the pGWB and ImpGWB series were kindly provided by Tsuyoshi Nakagawa (*22, 23*). pB7RWG2.0 and pBGYN are described in (*24*) and (*25*), respectively. The vectors used for CRISPR-Cas9-mediated genome editing are described in (*26*), and the sequences to target BAM1 and BAM2 were TCTCCGGTCTCAACCTCTC and CTCACCTCTCCTGACCTCA, respectively.

### Quantitative RT-PCR (qPCR)

Quantitative RT-PCR was performed as described in (*27*). Primers used are as follows: *C4*: ATCCGAACATTCAGGCAGCT and TGCTGACCTCCTCTAGCTGA (primer pair efficiency: 96%); *SlBAM1*: CTTGCCCTGAAAACTGCCAT and CGTGACACCATTCCACGTAC (primer pair efficiency: 90%). *ACTIN* (*ACT2*) was used as normalizer (*28*).

Quantitative PCR to determine viral accumulation was performed as described in (*27*), with primers to amplify *Rep*: TGAGAACGTCGTGTCTTCCG and TGACGTTGTACCACGCATCA (primer pair efficiency: 95%). 25S ribosomal DNA interspacer (ITS) was used as internal control (*29*).

### Quantification of *SUL* silencing

To quantify *SUL* silencing spread, pictures of leaves of the corresponding transgenic plants were transformed into black and white images (32 bit) and the area and black/white pixel ratio was calculated using ImageJ.

### Affinity-purification and mass spectrometry

Affinity-purification – mass spectrometry analysis (AP-MS) was performed as described in (*30*) with minor modifications. In order to efficiently extract membrane proteins, 2% of NP-40 was used in the protein extraction buffer.

### Yeast two-hybrid screen

Yeast two-hybrid screening was performed by Hybrigenics Services, S.A.S., Paris, France (http://www.hybrigenics-services.com). The coding sequence of C4 protein from *Tomato yellow leaf curl* virus was PCR-amplified and cloned into pB27 as a C-terminal fusion to LexA DNA-binding domain (LexA-C4). The construct was checked by sequencing the entire insert and used as a bait to screen a random-primed Tomato virus-infected plant cDNA library constructed into pP6. pB27 and pP6 derive from the original pBTM116 (*31*) and pGADGH (*32*) plasmids, respectively. 109 million clones (more than 10-fold the complexity of the library) were screened using a mating approach with YHGX13 (Y187 *ade2-101*::*loxP-kanMX-loxP*, *mat*α) and L40ΔGal4 (*mat*a) yeast strains as previously described (*33*). 318 His+ colonies were selected on a medium lacking tryptophan, leucine and histidine. The prey fragments of the positive clones were amplified by PCR and sequenced at their 5’ and 3’ junctions. The resulting sequences were used to identify the corresponding interacting proteins in the GenBank database (NCBI) using a fully automated procedure.

### Gel filtration assay

The C4 protein alone or co-expressed with the kinase domain of BAM1 was purified by Ni-affinity column followed by anion exchange column. The purified proteins were subjected to a gel filtration assay (Superdex 200 Increase 10/300 GL column; GE Healthcare).

### Viral infections

Viral infections in tomato were performed as described in (*27*).

### Northern blot of siRNA

Northern blots of siRNAs were performed as described in (*34*).

### Root elongation assays

Root elongation assays with CLV3p were performed as described in (*35*).

### Microprojectile bombardment assays

Microprojectile bombardment assays were performed as described (*36*). Four-to six-week-old expanded leaves of relevant Arabidopsis lines were bombarded with gold particles coated with pB7WG2.0.eGFP and pB7WG2.0.RFP_ER_ using a Bio-Rad Biolostic® PDS-1000/He Particle Delivery System. Bombardment sites were imaged 24 hours post-bombardment by epifluorescence (Leica DM6000) or confocal (Leica SP8) microscopy with a 25x water dipping lens and the number of cells to which GFP had spread was recorded (RFP_ER_ served as a marker to identify the transformed cell). For each line, data was collected from at least 3 independent bombardment events, each of which consisted of leaves from at least two individual plants. Statistical nonparametric Mann-Whitney analysis was performed using Genstat software.

## SUPPLEMENTARY FIGURES

**Figure S1.**
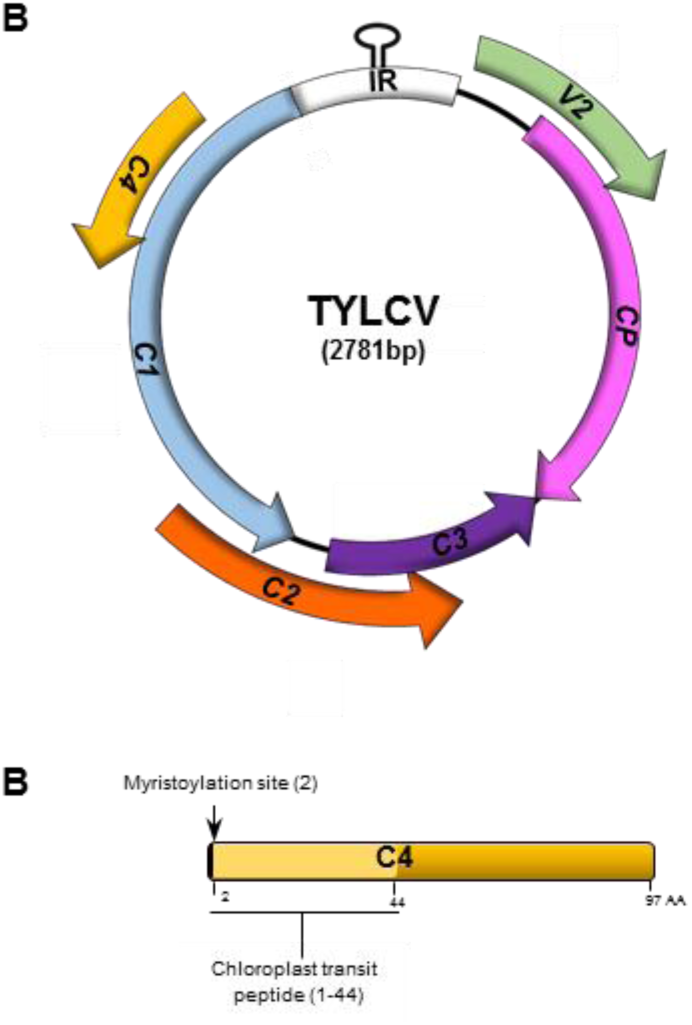
Genomic organization of *Tomato yellow leaf curl virus* (TYLCV) (A) and localization motifs in its C4 protein (B).

**Figure S2.**
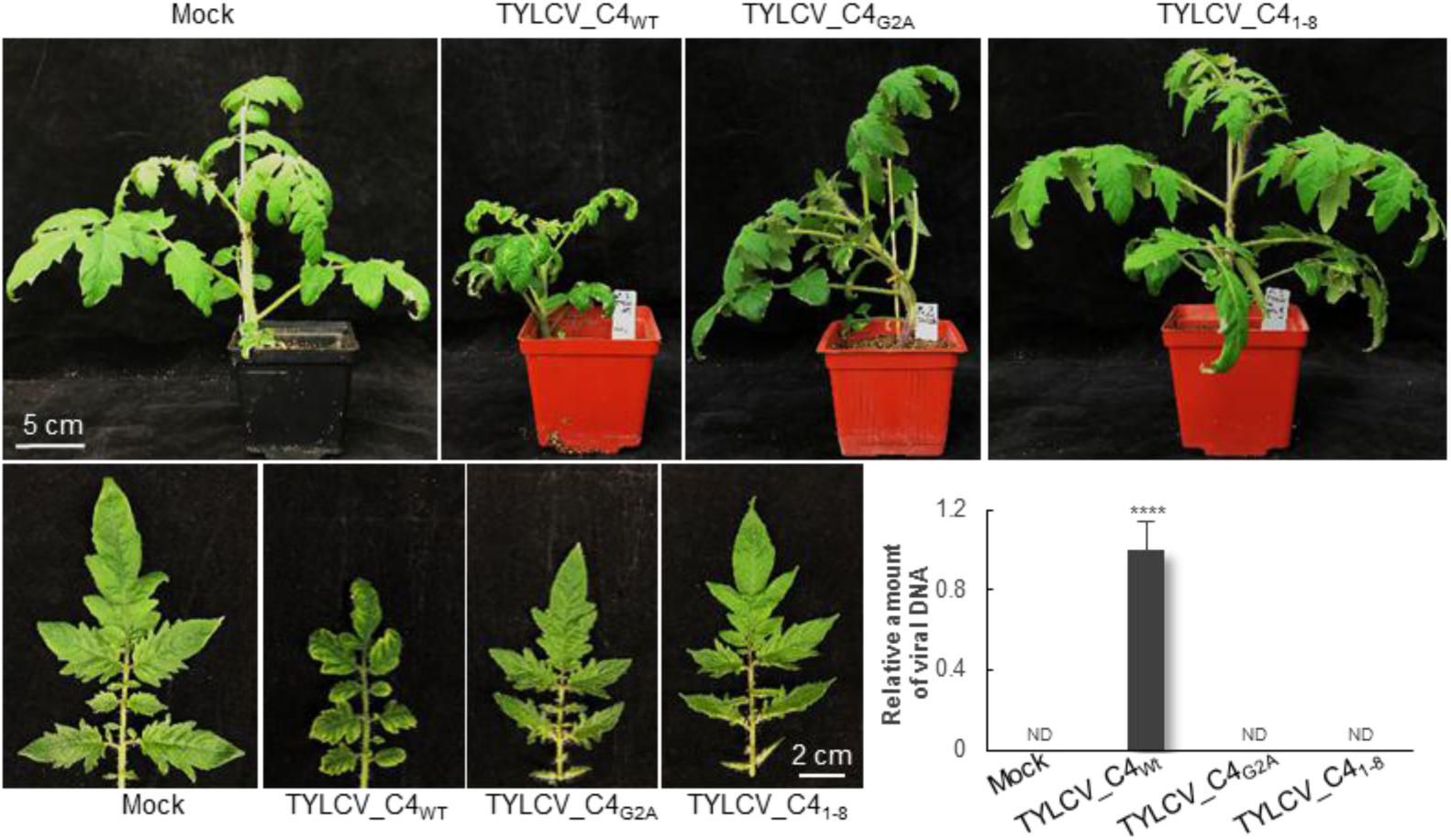
Tomato plants infected with TYLCV wild type, o mutant versions to express a non-myristoylable C4 (TYLCV_C4_G2A_), or the first eight amino acids of C4 only (TYLCV_C4_1-8_). The graph represents the relative viral DNA accumulation in plants inoculated with TYLCV variants or mock as determined by real-time PCR of total DNA extracted from apical leaves at 21 dpi. Values represent the average of 4 plants, and are relative to that of TYLCV_C4_WT_. Bars represent SE. Asterisks indicate a statistically significant difference (****, p-value < 0.0001), according to a Student’s t-test; ND: not detectable.

**Figure S3.**
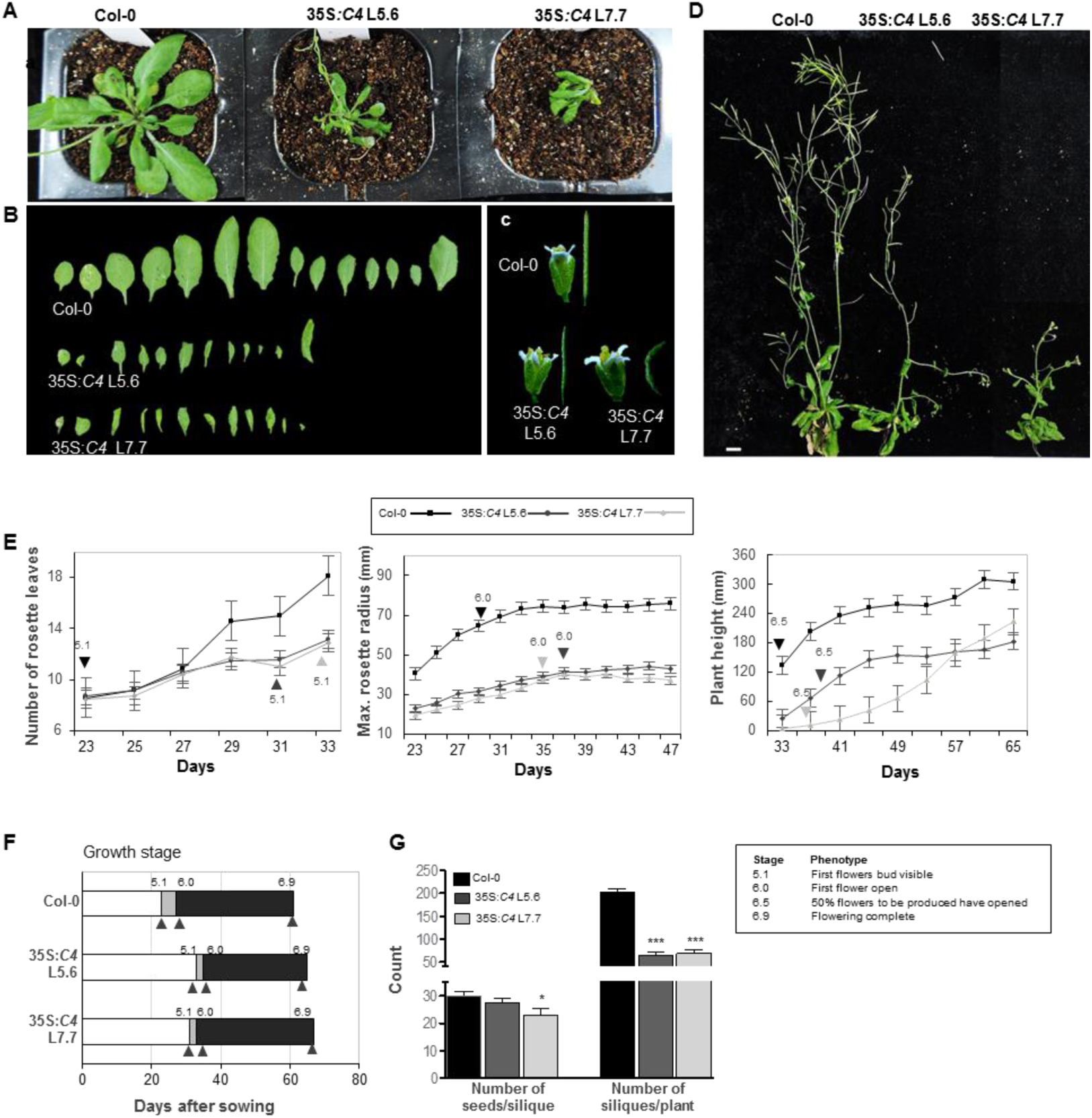
Developmental phenotype of transgenic Arabidopsis plants expressing C4. A. Five-week-old transgenic plants grown in long day conditions. B. Leaves of five-week-old plants grown in long day conditions. C. Representative flowers and siliques. D. Flowering seven-week-old plants. E. Number of rosette leaves, maximum rosette radius, and plant height. Results are the average of ten plants. Bars represent SE. F. Growth stage progression. Results are the average of ten plants. G. Numbers of seeds/silique and of siliques/plant. Results are the average of six plants.

**Figure S4.**
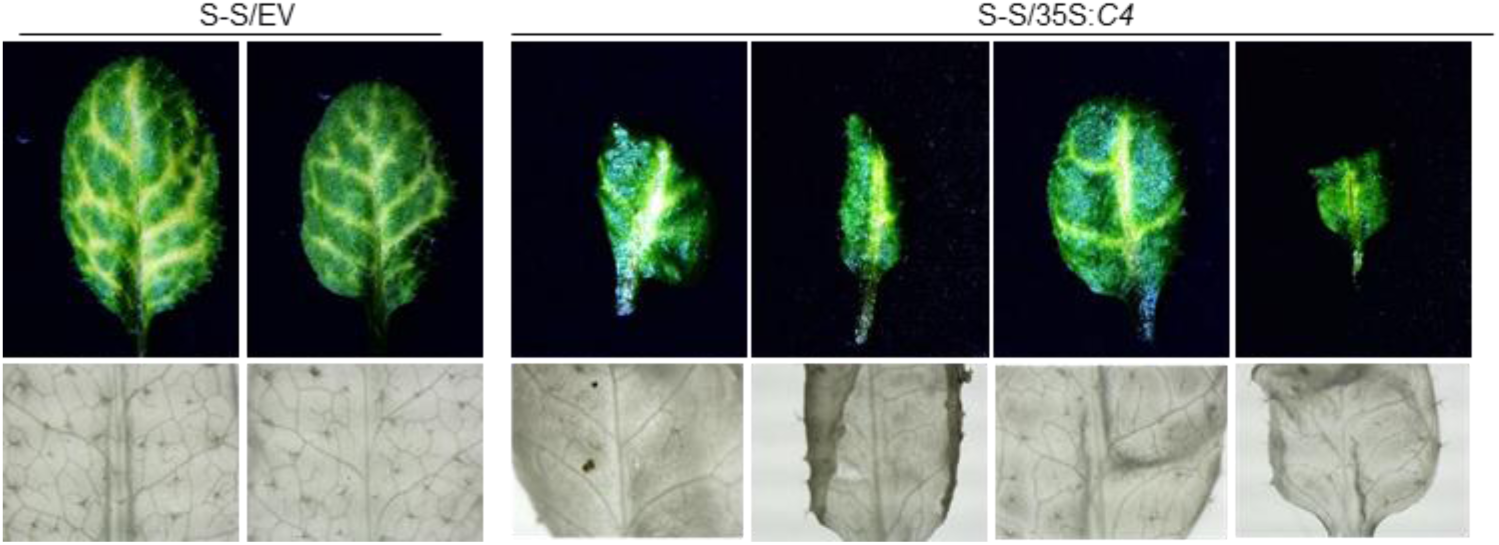
Venation pattern in the C4-expressing SUC:*SUL* plants (S-S/35S:*C4*) and the SUC:*SUL* control plants (S-S/EV). Each leaf comes from one T1 plant.

**Figure S5.**
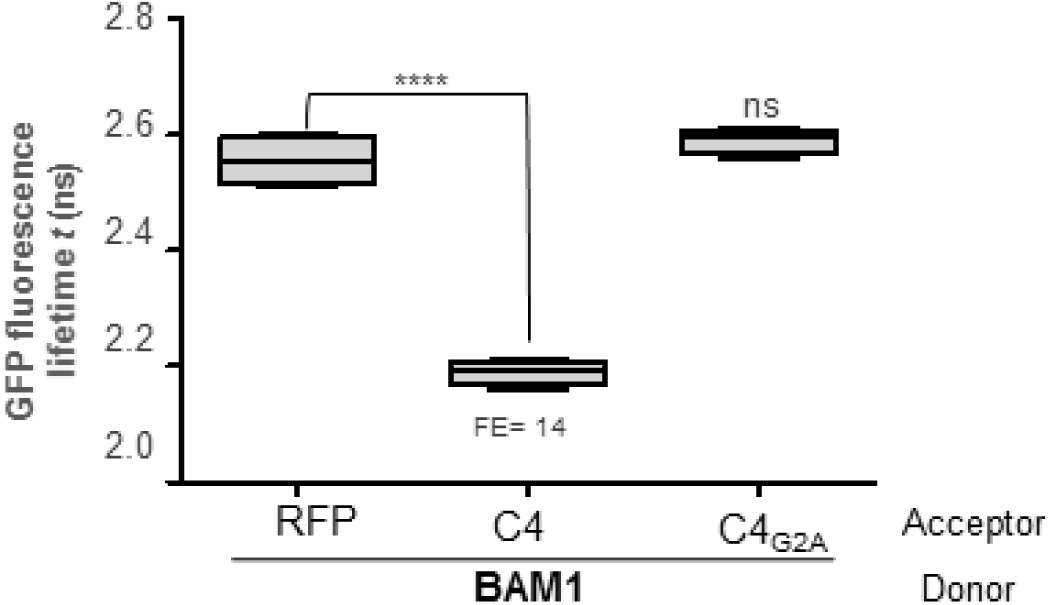
C4_G2A_ does not interact with BAM1 by FRET-FLIM upon transient co-infiltration in *N. benthamiana* leaves. Asterisks indicate a statistically significant difference (****, p-value < 0.0001), according to a Student’s t-test; ns: not significant.

**Figure S6.**
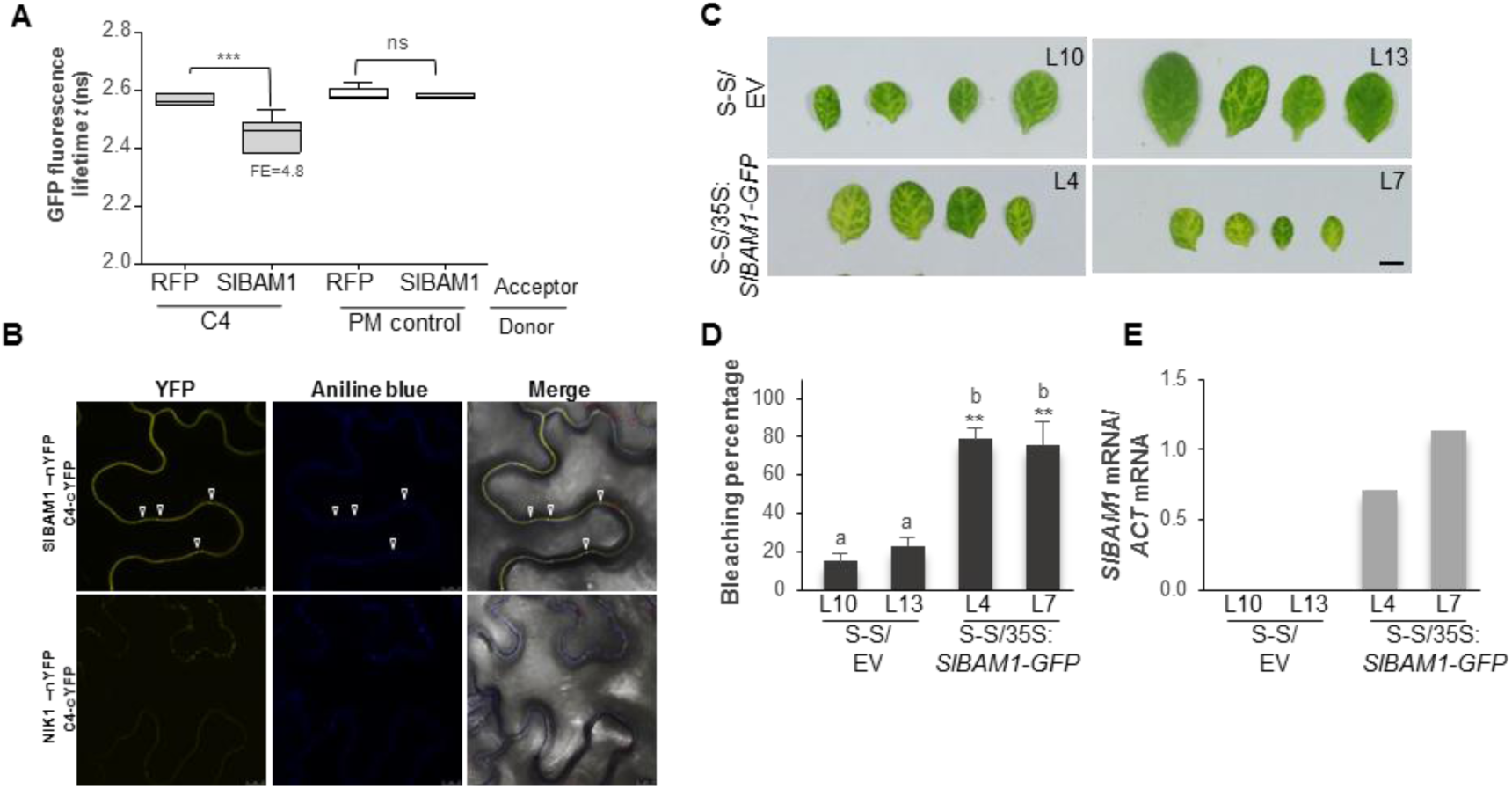
C4 interacts with BAM1 from tomato (SlBAM1). A. Interaction between C4 and SlBAM1 by FRET-FLIM upon transient co-infiltration in *N. benthamiana* leaves. The membrane protein NP_564431 (NCBI) is used as a negative control (PM control). FE: FRET efficiency. Asterisks indicate a statistically significant difference (***, p-value<0.005), according to a Student’s t-test; ns: not significant. B. Interaction between C4 and SlBAM1 by BiFC upon transient co-infiltration in *N. benthamiana* leaves. The receptor-like kinase NIK1 is used as a negative control. Arrowheads indicate plasmodesmata. C. Leaves of four-week-old transgenic SUC:*SUL* plants overexpressing SlBAM1-GFP (S-S/35S:*SlBAM1-GFP*) or transformed with the empty vector (S-S/EV). Each set of leaves comes from one T2 plant from an independent line. D. Quantification of the bleaching percentage of the leaves in (C). Scale bar: 0.5 cm. Bars represent SE; bars with the same letters are not significantly different (P=0.05) according to Dunnet’s multiple comparison test. E. Expression of *SlBAM1* in transgenic S-S/35S:*SlBAM1-GFP* relative to actin mRNA, as measured by qPCR.

**Figure S7.**
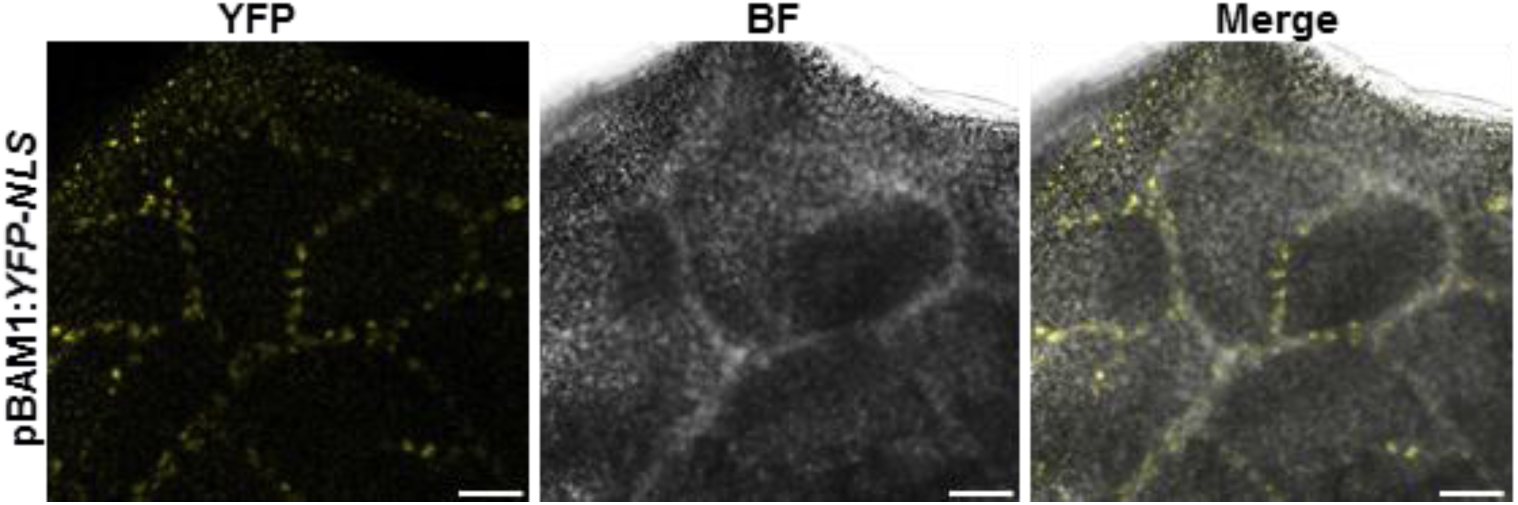
*BAM1* is strongly expressed in the vasculature. Leaves of twelve-day-old transgenic pBAM1:*YFP-NLS* Arabidopsis plants. BF: Bright field. Scale bar: 50 μm.

**Figure S8.**
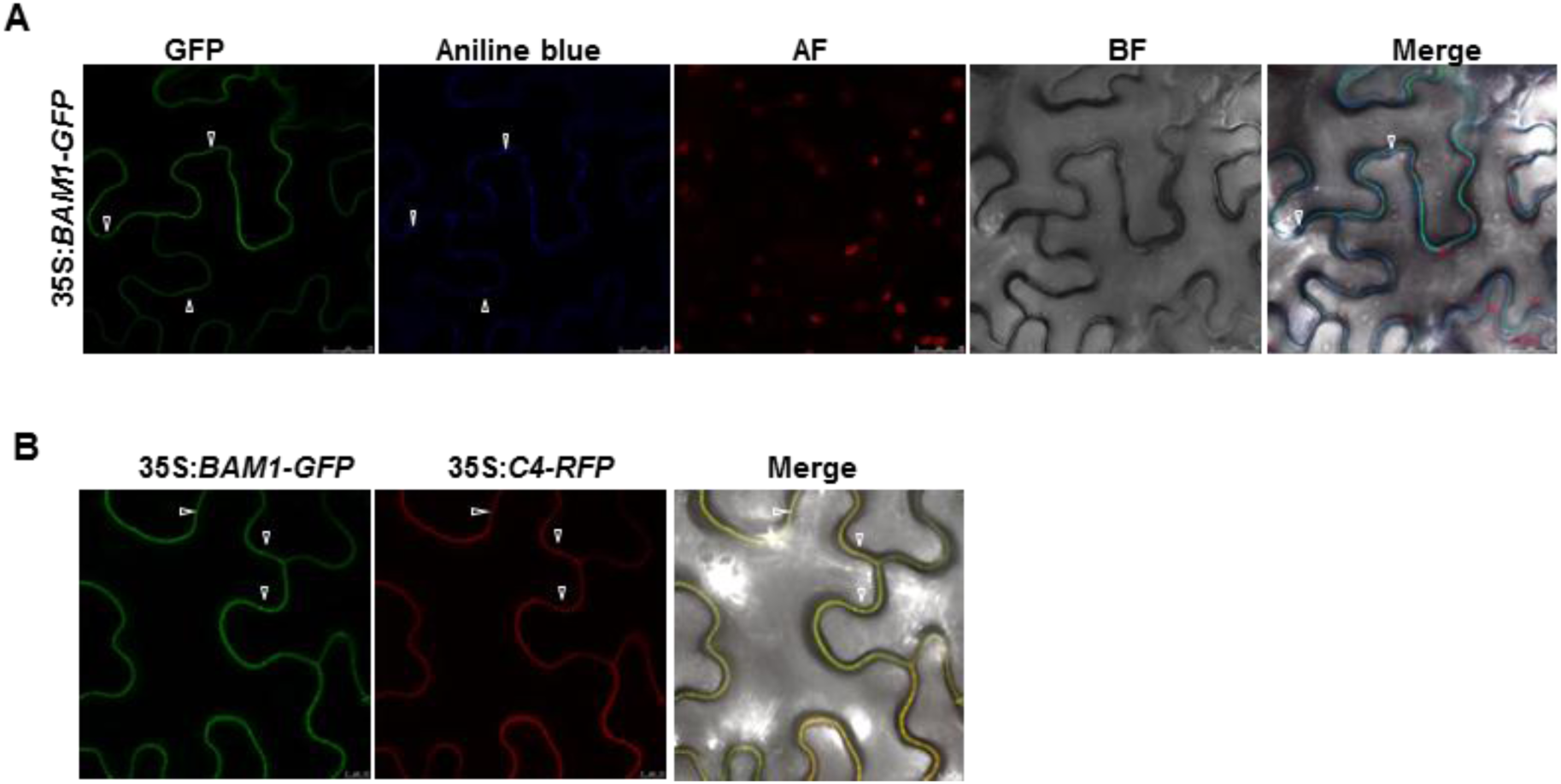
BAM1 co-localizes with C4 at plasma membrane and plasmodesmata. A. Subcellular localization of BAM1-GFP upon transient expression in *N. benthamiana* leaves. AF: Autofluorescence. BF: Bright field. B. Subcellular co-localization of BAM1-GFP and C4-RFP upon transient co-expression in *N. benthamiana* leaves. Arrowheads indicate plasmodesmata.

**Figure S9.**
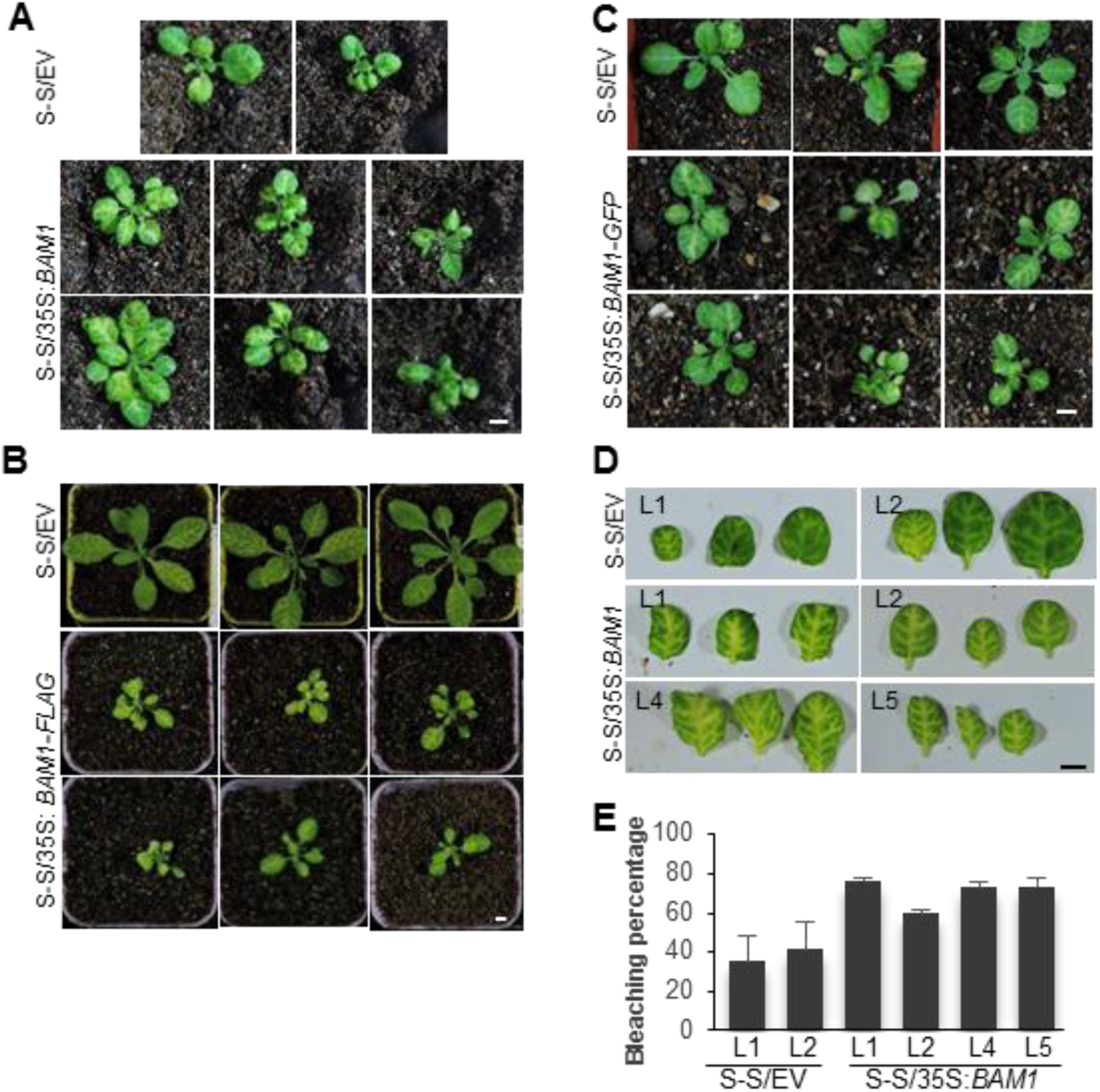
Phenotype of SUC:*SUL* plants overexpressing BAM1, BAM1-GFP, or BAM1-FLAG. A. Transgenic SUC:*SUL* lines overexpressing BAM1 (S-S/35S:*BAM1*) or transformed with the empty vector (S-S/EV). B. Transgenic SUC:*SUL* lines overexpressing BAM1-FLAG (S-S/35S:*BAM1-FLAG*) or transformed with the empty vector (S-S/EV). C. Transgenic SUC:*SUL* lines overexpressing BAM1-GFP (S-S/35S:*BAM1-GFP*) or transformed with the empty vector (S-S/EV). D. Leaves of four-week-old transgenic SUC:*SUL* plants overexpressing BAM1 (S-S/35S:*BAM1*) or transformed with the empty vector (S-S/EV). Each set of leaves comes from one T1 plant. E. Quantification of the bleaching percentage of the leaves in (D). Bars represent SE. Scale bar: 0.5 cm.

**Figure S10.**
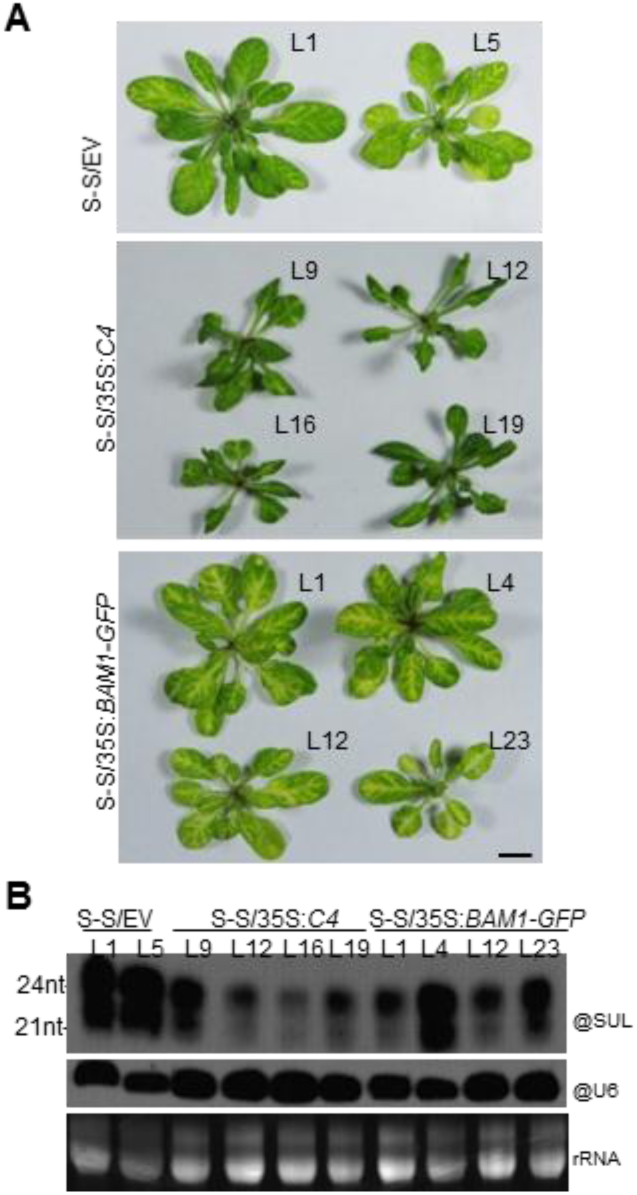
Phenotype (A) and *SUL* siRNA accumulation (B) of SUC:*SUL*/35S:*C4* (S-S/35S:*C4*), SUC:*SUL*/35S:*BAM1-GFP* (S-S/*BAM1-GFP*) lines, or SUC:*SUL* plants transformed with the empty vector (S-S/EV). Each plant comes from one T2 independent line.

**Figure S11.**
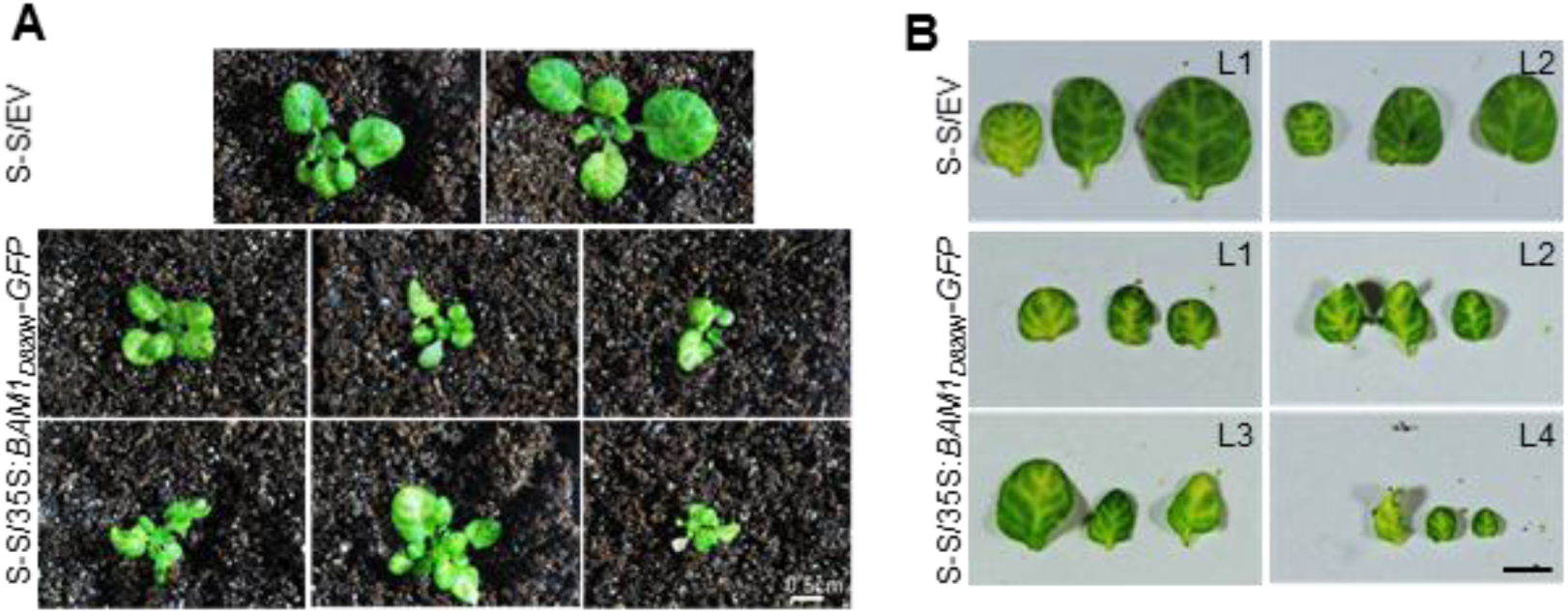
Phenotype of SUC:*SUL*/35S:*BAM1*_*D820N*_*-GFP* lines. A. Four-week-old T1 plants grown in long-day conditions. B. Leaves of representative plants from (A). Scale bar: 0.5 cm.

**Figure S12.**
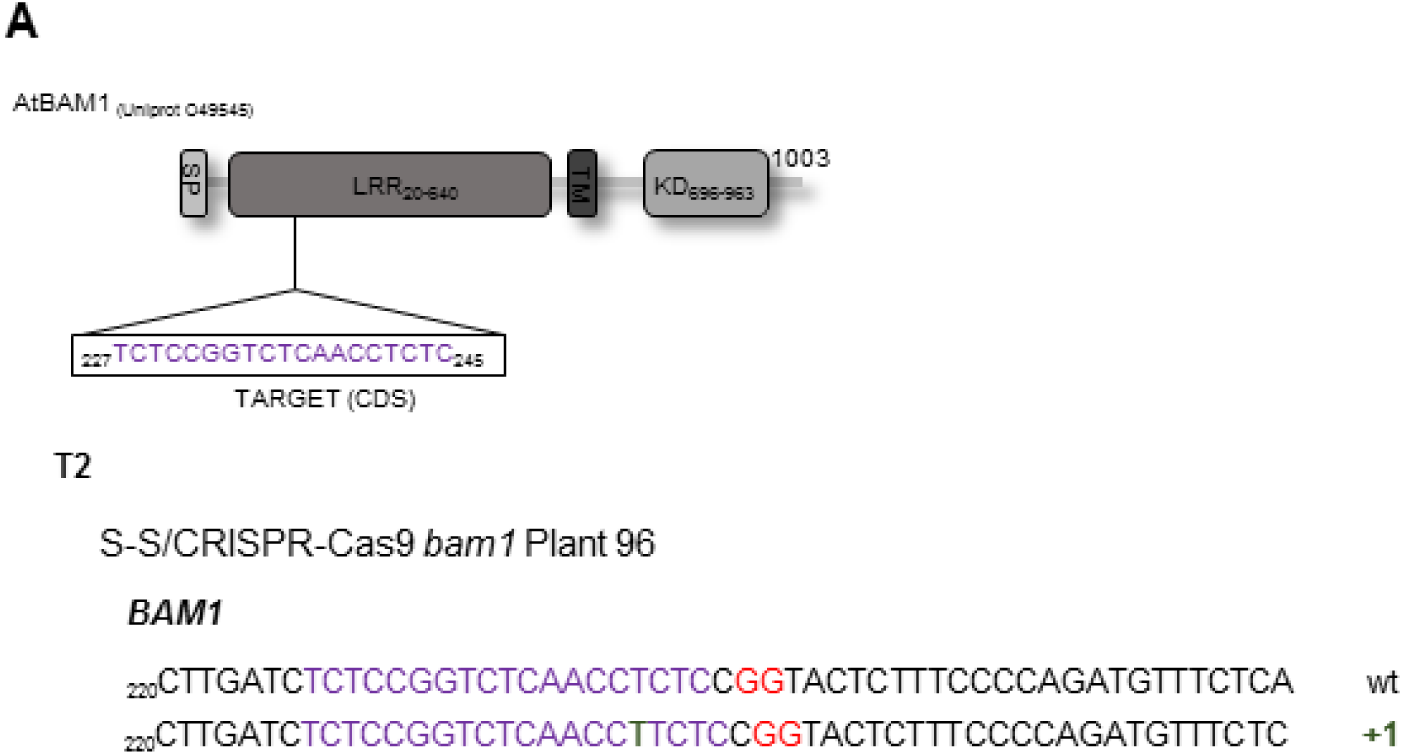
Details of the *bam1* mutations in the *bam1* single mutant generated by CRISPR-Cas9. The target of the CRISPR-Cas9 construct and the mutation in the plant depicted in Figure 3 are indicated (for details, see Table S1).

**Figure S13.**
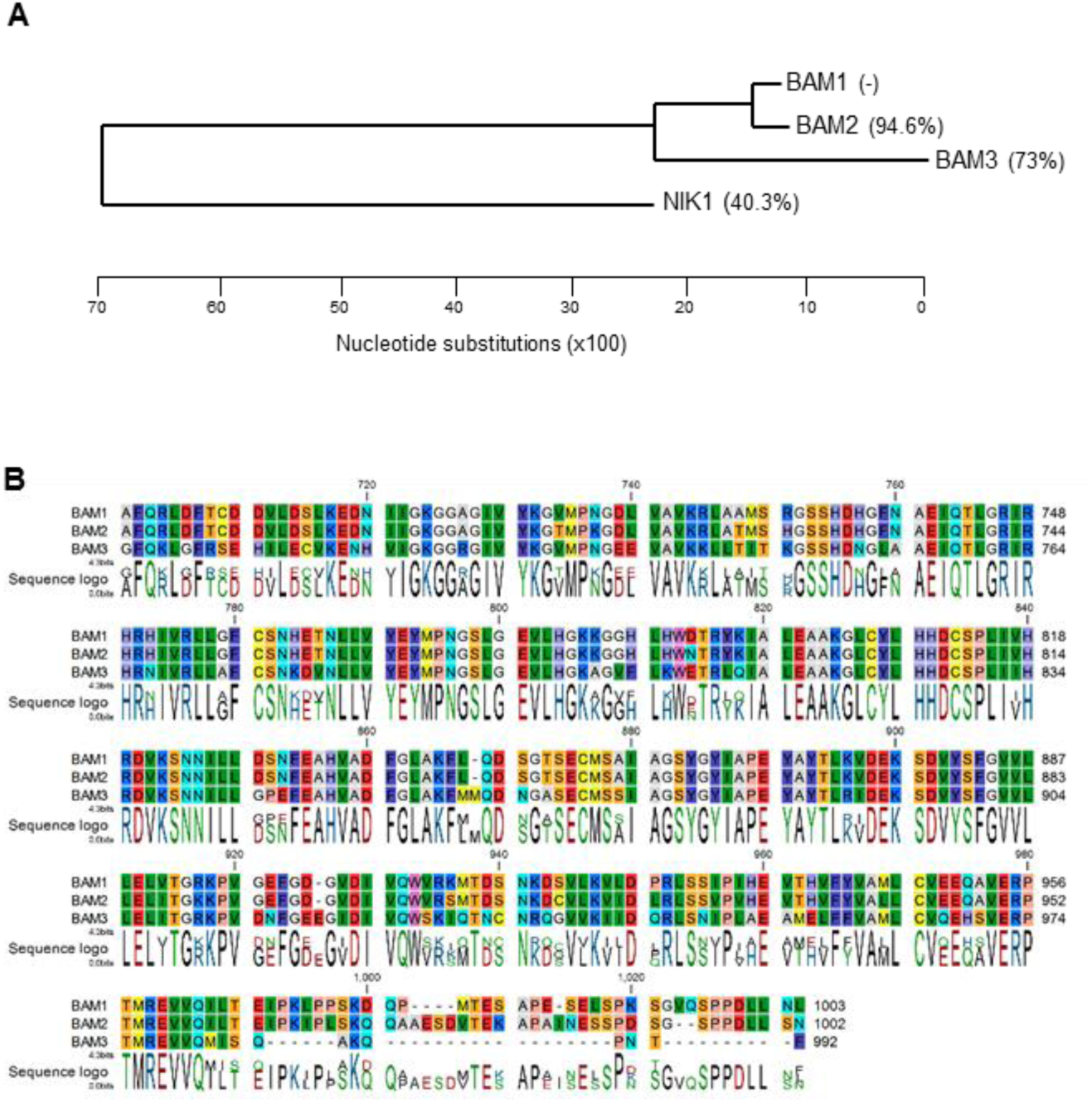
Phylogeny of the BAM1 family of receptor-like kinases. A. Phylogenetic tree of BAM1, BAM2, and BAM3, based on the kinase domain. The unrelated receptor-like kinase NIK1 is used as outgroup. Percentage of identity to BAM1 is indicated in parenthesis. B. Alignment of the kinase domain of BAM1, BAM2, and BAM3. The phylogenetic analysis and sequence alignment were performed using CLC Main Workbench 7.

**Figure S14.**
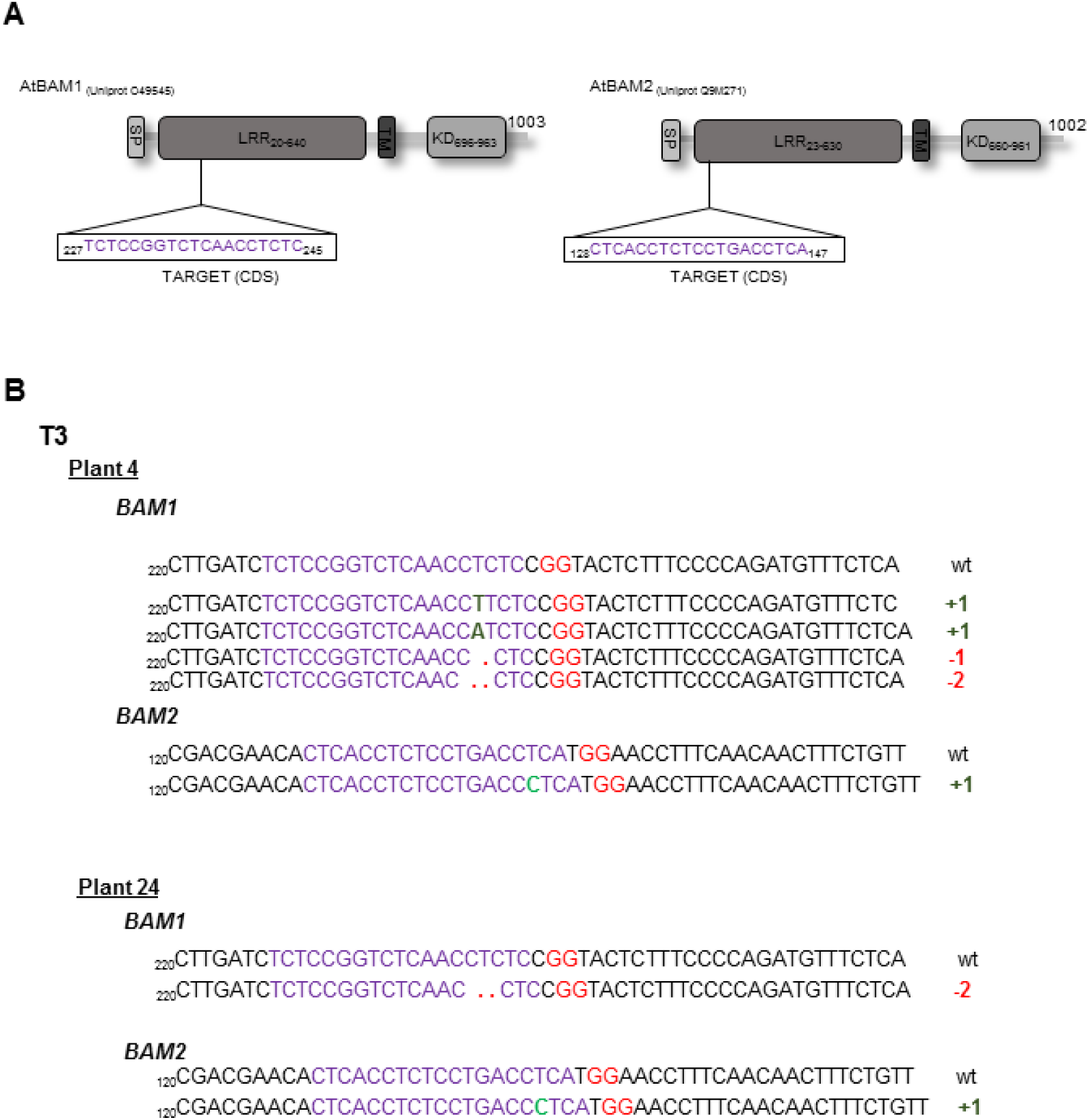
Details of the *bam1* and *bam2* mutations in the *bam1 bam2* double mutants generated by CRISPR-Cas9. The target of the CRISPR-Cas9 construct and the mutations in the plants depicted in Figure 4 are indicated (for details, see Table S1).

**Figure S15.**
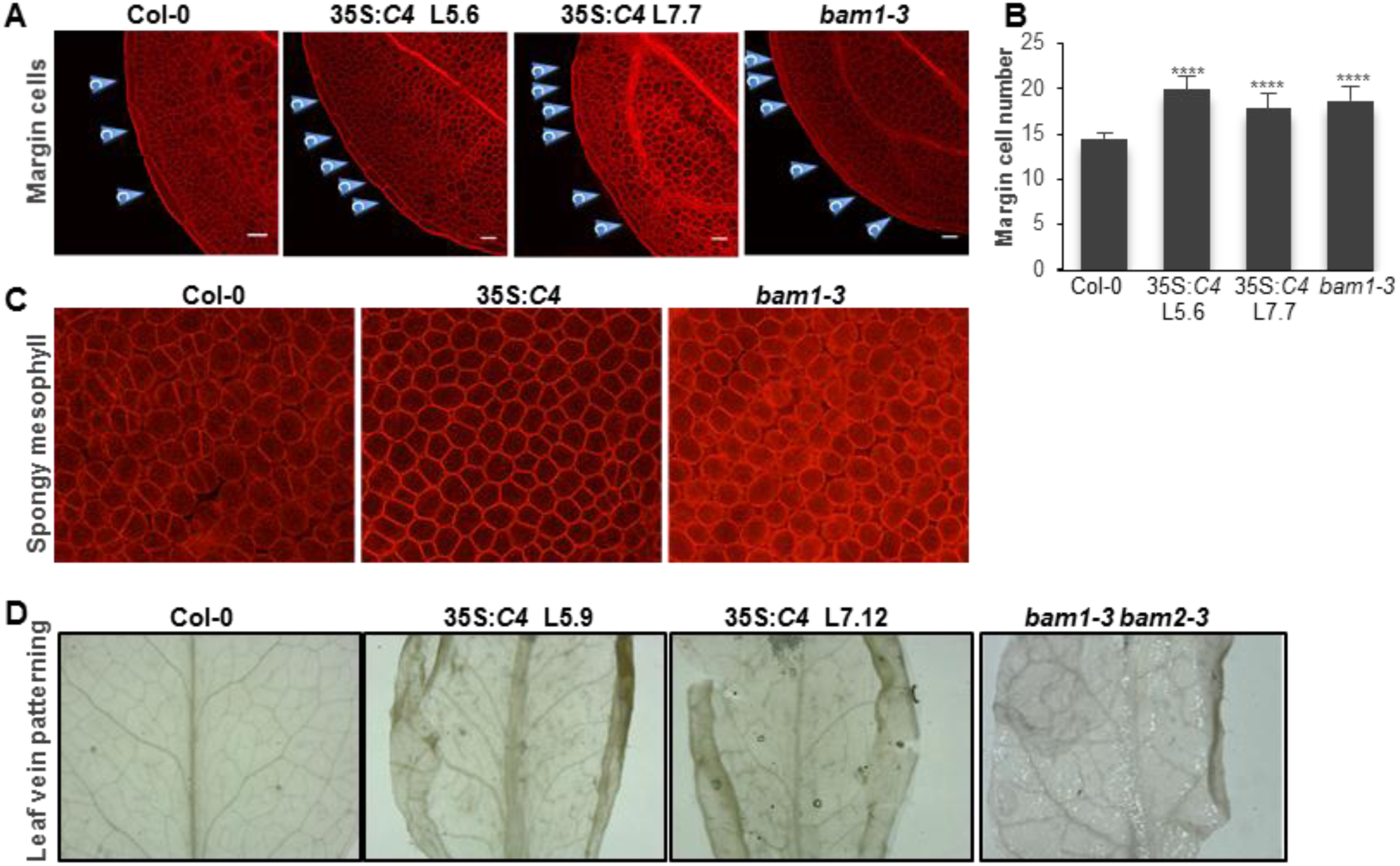
Similar developmental phenotypes in the transgenic Arabidopsis lines expressing C4 and the *bam1-3* single or *bam1-3 bam2-3* double mutant. A. Margin cells in cotyledons of two-day-old seedlings. Asterisks indicate a statistically significant difference (****, p-value <0.0001), according to a Student’s t-test. B. Margin cell numbers of cotyledons of two-day-old seedlings. C. Spongy mesophyll of cotyledons of two-day-old seedlings. D. Leaf vein patterning in leaves of five-week-old plants.

**Figure S16.**
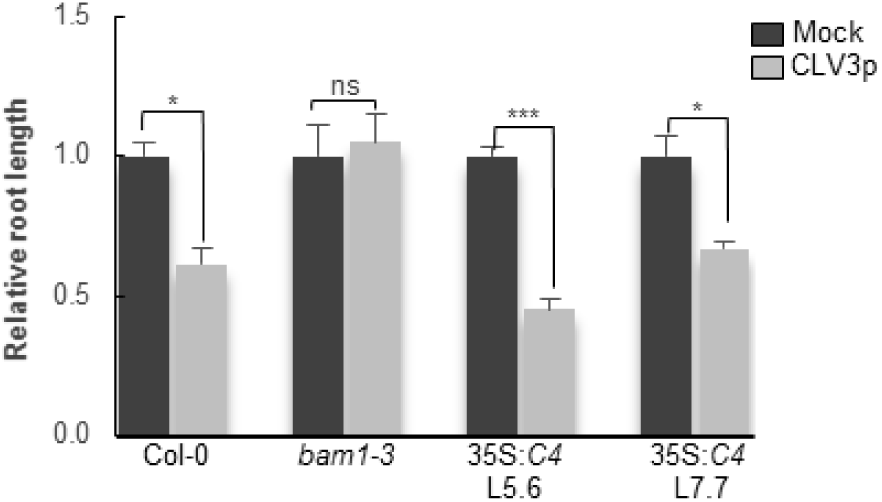
Root response to CLV3p in transgenic seedlings expressing C4. Wild-type (Col-0) and *bam1-3* mutant seedlings are used as control. Asterisks indicate a statistically significant difference (***,p-value < 0.005; *, p-value < 0.05), according to a Student’s t-test; ns: not significant.

**Figure S17.**
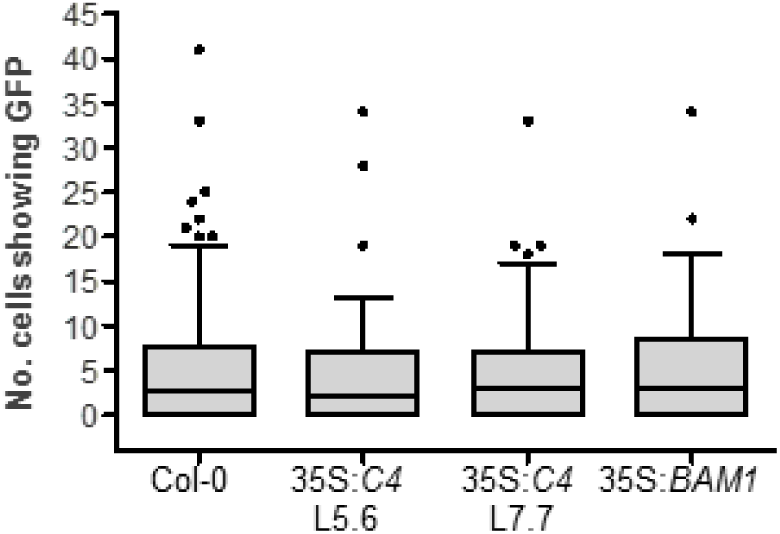
Plasmodesmal flux is not altered in lines that overexpress C4 and BAM1. Box plot of data from microprojectile bombardment assays. 35S:*GFP* diffused to the same number of cells in wild-type Col-0 and the overexpression lines 35S:*C4* L5.6 (p-value 0.658), 35S: *C4* L7.7 (p-value 0.831) and 35S:*BAM1* (p-value 0.926). Boxes indicate the interquartile range with the line indicating the median value; whiskers are 1.5 time the interquartile range.

**Table S1.**
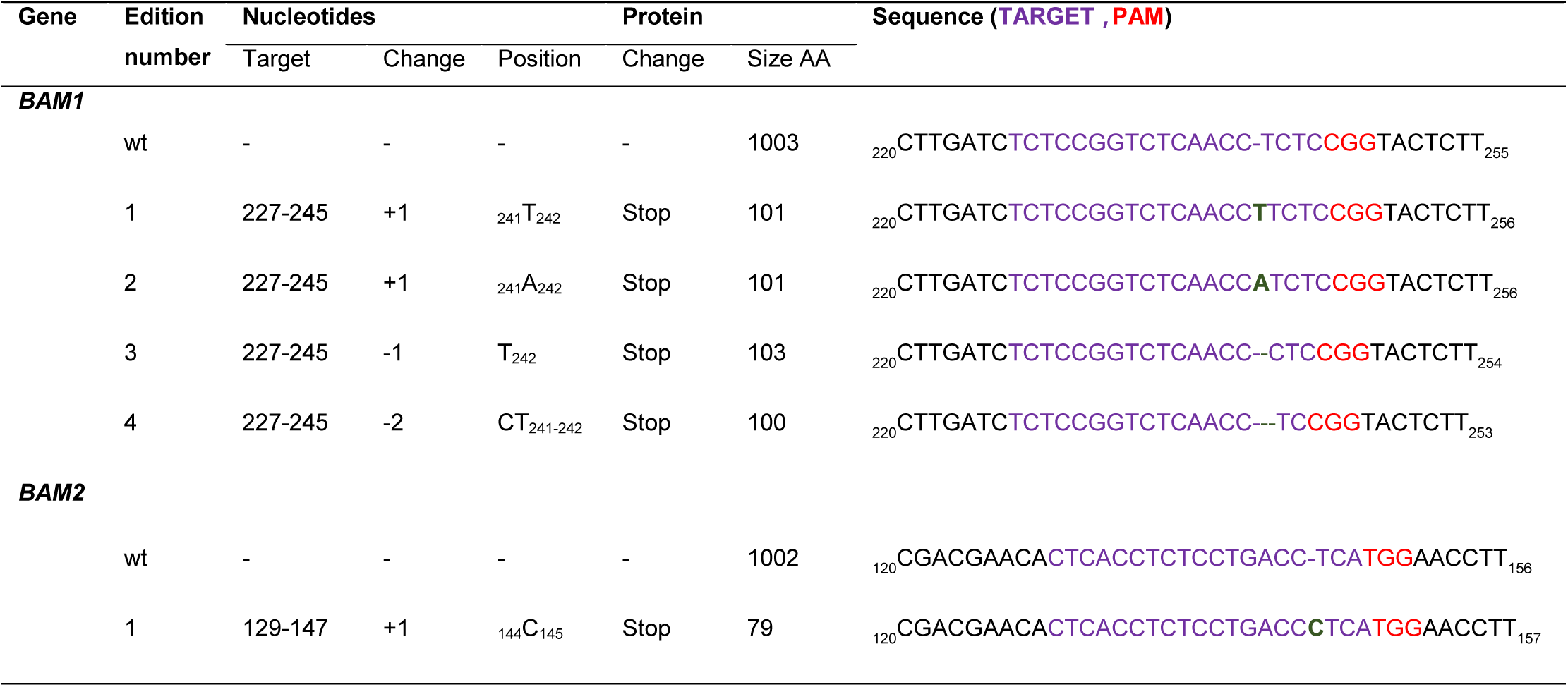
Mutations in *BAM1* and *BAM2* obtained by CRISPR-Cas9

| Gene | Generation | Number | Editions |  |
| --- | --- | --- | --- | --- |
|  |  |  | *BAM1* | *BAM2* |
| *BAM1* |  |  |  |  |
|  | T2 | Plant 96 | Edition 1-/- | wt |
|  | T3 | Plant 4 | Edition 1, Edition 2, Edition 3, Edition 4 | Edition 1-/- |
|  | T3 | Plant 24 | Edition 4-/- | Edition 1-/- |

**Table S2.**
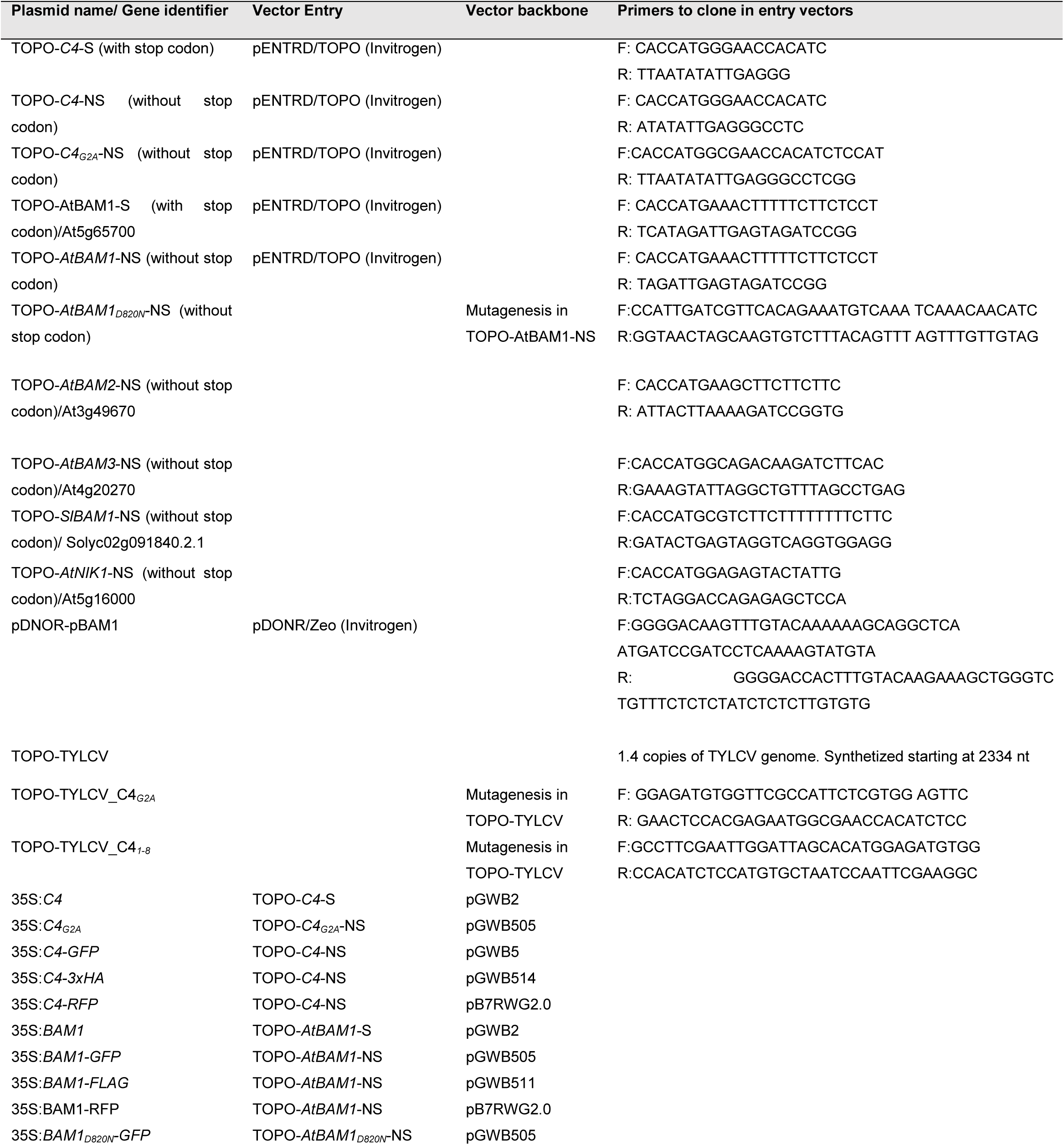
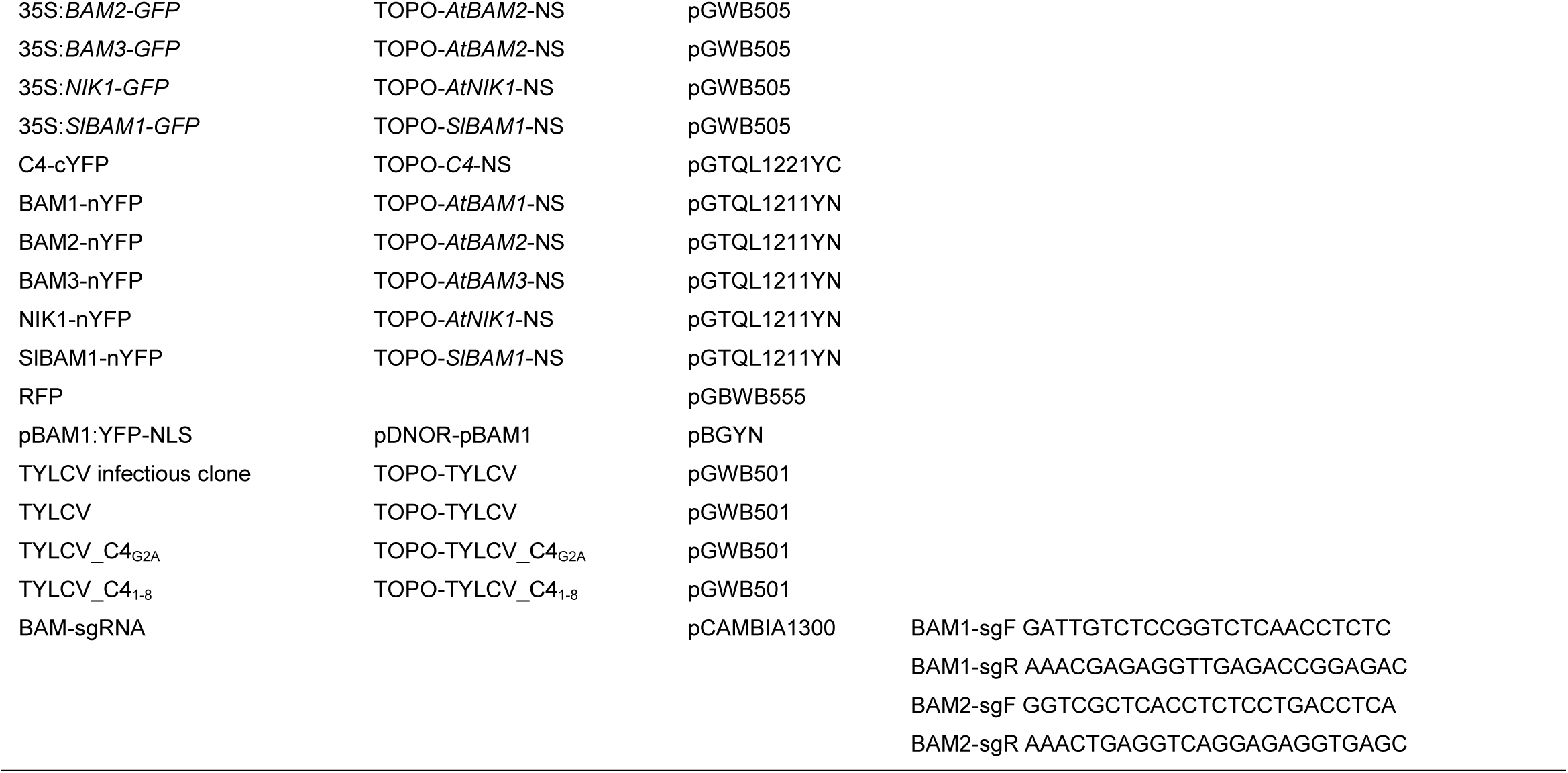
Plasmids generated in this work.

